# Multilevel analysis of response to plant growth promoting and pathogenic bacteria in Arabidopsis roots and the role of CYP71A27 in this response

**DOI:** 10.1101/2025.03.26.645393

**Authors:** Anna Koprivova, Daniela Ristova, Miroslav Berka, Veronika Berková, Gözde Merve Türksoy, Tonni Grube Andersen, Philipp Westhoff, Martin Černý, Stanislav Kopriva

## Abstract

Understanding how plants distinguish between commensal and pathogenic microorganisms is one of the major challenges in the plant microbe interaction research. We previously identified a gene encoding CYP71A27 connected to camalexin, which is necessary for a plant growth promoting (PGP) activity of a number of bacterial strains. To dissect its function, we compared multilevel responses in roots of wild type Arabidopsis and the *cyp71A27* mutant to two bacterial strains, a PGP *Pseudomonas fluorescens* CH267 and a pathogen *Burkholderia glumeae* PG1. We show that incubation with these bacteria leads to significant and distinct transcriptional reprogramming. This is accompanied by a proteome remodelling in both shoots and roots and profound changes in accumulation of many metabolites, primarily sugars, amino acids, and TCA cycle intermediates, but also by alterations in the ionome. We then analysed the mutant *cyp71a27* and identified number of genes and proteins differently regulated, particularly after interaction with the PGP strain, but only a very mild impact of the mutation on root metabolites and exudates. We analysed a variety of mutants in genes differentially regulated by *Pseudomonas* sp. CH267 in *cyp71a27* and revealed that their response to this PGP bacterial strain is similar to *cyp71a27*. Thus, it seems that CYP71A27 is a non-canonical P-450 without a metabolic function but active in signalling pointing to a regulatory role of the CYP71A27 gene, particularly in interaction with a plant growth promoting bacteria.

## INTRODUCTION

Plants in their natural habitats interact with a great variety of microorganisms, some commensal, some harmful. To protect against the latter, plants developed a complex multi-layered innate immunity system, including a chemical defense by diverse phytoalexins. However, both beneficial and pathogenic microbes are recognised by the same pattern recognition receptors through conserved microbial structures, therefore, a central question is, how they are distinguished by the immune system (Thoms et al., 2021; Ji et al., 2022). Indeed, even the symbiotic mycorrhizal fungi and Rhizobia initially trigger an immune response (Campos-Soriano et al., 2012), but are able to overcome it, e.g., by bacterial polysaccharides chelating calcium ions (Aslam et al., 2008) or various effectors (de Jonge et al., 2010). The same is true for the plant growth promoting (PGP) bacteria which in plants induce genes related to immunity (Teixeira et al., 2021). While large number of studies elucidated the effects of PGP on the plants and different mechanisms of action have been identified, including facilitating the supply of nutrients, synthesis of plant hormones, or protection against pathogens (Kertesz and Mirleau, 2004; Poupin et al., 2013; Haney et al., 2015), how these strains overcome the plant immune system is still largely elusive. It seems, however, that metabolic signalling is at the core of the symbiosis establishment, either through selective effects of plant metabolites on specific microbes or reciprocal signalling through the symbionts (Zipfel and Oldroyd, 2017; Sasse et al., 2018; Huang et al., 2019; Harbort et al., 2020; Thoms et al., 2021). Indeed, for example, mutualistic interactions of Arabidopsis with endophytic fungi are dependent on fungal growth restriction by indolic glucosinolates and other indole-derived phytoalexins (Lahrmann et al., 2015). Accordingly, many plant metabolites with antimicrobial properties have been identified as compounds shaping the composition of plant microbiome (reviewed in (Jacoby et al., 2021).

One of these metabolites is camalexin, a sulfur-containing indolic compound found in the Brassicaceae (Glawischnig, 2007; Koprivova et al., 2019). Camalexin has been characterized as a phytoalexin induced by fungal and bacterial infection, but also by abiotic factors (Browne et al., 1991; Kliebenstein et al., 2005; Nguyen et al., 2022). It is an important component of the immunity against fungal pathogens, such as *Botrytis cinerea*, but it is also exuded in the rhizosphere upon elicitation with flagellin (Rowe and Kliebenstein, 2008; Millet et al., 2010). Camalexin was found to modulate microbial sulfatase activity in the soil, specifically, a variation in a gene from camalexin biosynthetic network, *CYP71A27*, was identified in a genome-wide associated analysis of variation in sulfatase activity among Arabidopsis accessions (Koprivova et al., 2019). Also other mutants in camalexin synthesis genes *cyp71A12* and *cyp71A13* possessed lower camalexin concentration in the roots and stimulated less sulfatase activity in the rhizosphere. Additionally, the *cyp71A27* mutant was not able to profit from interaction with plant growth promoting (PGP) bacteria *Pseudomonas* sp. CH267 or *P. simiae* WCS417r (Koprivova et al., 2019). Thus, camalexin has a dual function as phytoalexin in defense against diverse fungal pathogens and as a metabolite important for microbiome function and PGP activity of various bacterial strains (Kliebenstein et al., 2005; Koprivova et al., 2019). The mechanism of camalexin action and the molecular function of the CYP71A27, are however not known yet.

We found that the CYP71A27 is inactive in camalexin synthesis and thus addressed its function by multiomics analyses of the *cyp71A27* mutant in roots response to two bacterial strains, a PGP strain *Pseudomonas* sp. CH267 and a pathogen *B. glumae* PG1. We show that incubation with these bacteria leads to significant transcriptional reprogramming and different accumulation of a range of metabolites and that CYP71A27 is particularly important for interaction with the PGP strain. Analysis of mutants in several genes differentially regulated by *Pseudomonas* sp. CH267 in *cyp71a27* revealed differences in response to this PGP bacterial strain similar to the *cyp71a27*, pointing to a regulatory role of the *CYP71A27* gene in the interaction of Arabidopsis with PGP bacteria.

## RESULTS (4624 words)

### CYP71A27 is not active in camalexin synthesis

As a first step in deciphering the function of CYP71A27 we confirmed its localization. Promoter-reporter study indicated that the gene is expressed in the vasculature (Koprivova et al., 2019). When applying a Clearsee staining protocol (Ursache et al., 2018) on 8-day-old roots of a plant line containing a vector that allowed CYP71A27::GFP fusion protein expression from the native promoter, we found signal specifically localised in the phloem companion cells of the roots (Figure 1A, B). In addition, the fluorescence pattern appeared in a reticulate pattern, which suggested that the protein in localized in the endoplasmic reticulum, as expected for a cytochrome P-450 (Figure 1B). CYP71A27 has been described as a part of camalexin synthesis network and its loss leads to reduced camalexin levels (Koprivova et al., 2019). This can be caused by deficient camalexin production in the mutant or by direct signalling function of the protein independent from its enzymatic activity. To test whether CYP71A27 is involved in camalexin synthesis, its ability to complement the biosynthetic mutant *cyp71A12 cyp71A13* (Muller et al., 2015) was determined. This mutant accumulates only very low quantities of camalexin after treatment with *B. glumae* (Figure 1C, D). Complementation of the mutant with *35S::CYP71A12* as a control resulted in restoration of the camalexin synthesis after the pathogen treatment (Figure 1C). On the other hand, expression of CYP71A27 under the control of 35S promoter did not increase camalexin production, although the transcript was present to much higher levels than in WT (Figure 8D). Therefore, the CYP71A27 is likely not involved in camalexin synthesis and its effects on camalexin and transcriptional responses to bacteria are likely due to an indirect, possibly signal-related function, which may transverse the phloem tissue.

**Figure 1.**
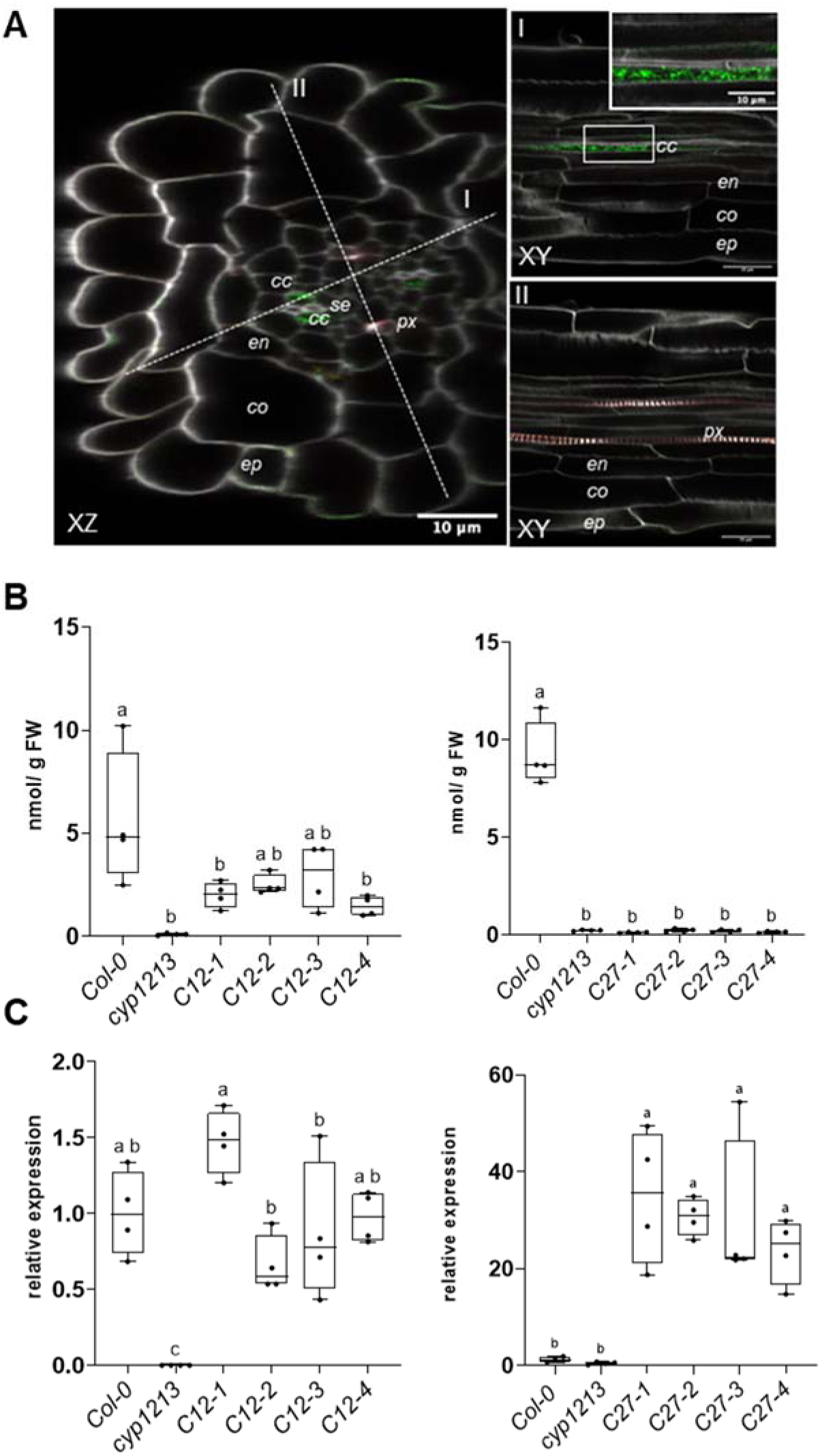
Characterization of CYP71A27. **A** Root sections of 8 days old Arabidopsis plants expressing a CYP71A27::GFP fusion protein (green) under the control of its native promoter region. Lignin was stained with basic fuchsin (red) and cellulose with Calcofluor white (grey). co: Cortex, cc: Companion cell, ep: Epidermis, en: Endodermis, px: Protoxylem, se: Sieve element. **B, C** The *cyp71A12 cyp71A13* mutant was complemented with *CYP71A12* or *CYP71A27* expressed under control of 35S promoter. WT, *cyp71A12 cyp71A13 (cyp1213)* and four independent complemented lines per construct were treated with *B. glumae*. **B** Camalexin levels were determined in the shoots. **C** Relative transcript levels of the transgenes were determined in the roots. The qRT-PCR results were obtained from 4 biological and 2 technical replicates, the transcript levels in WT Col-0 were set to 1. Letters mark values significantly different at P<0.05 (ANOVA).

### Transcriptional response of cyp71A27 and WT Arabidopsis roots to pathogenic and PGP bacterial strains

To identify the signalling role of CYP71A27 in responses to PGP and pathogen bacteria we analysed how the transcriptome responds in roots of the *cyp71a27* mutant and wild type (WT) Col-0 Arabidopsis roots when challenged with *Pseudomonas* sp. CH267 (further denoted as CH267) or *B. glumae* PG1, respectively. After 3 days co-inoculation with CH267, we found 862 differentially regulated genes (DEGs) (q<0.01, |log_2_FC|>1) in WT, while 1326 DEGs were found in *cyp71A27* (Figure 2A; Supplemental Table S1). The treatment with *B. glumae* resulted in a more substantial transcriptional reprogramming (Figure 2B; Supplemental Table S1), however, the response to the two bacterial strains partially overlapped. A principal component analysis (PCA) revealed that only after treatment with CH267 a significant separation of the *cyp71A27* and the WT was observed in the first two components (Figure 2C). Among the DEGs upregulated by CH267, 62% and 48% also responded in *B. glumae* -treated WT and *cyp71A27*, respectively, while among the downregulated DEGs these constituted 80% and 58% respectively (Supplemental Figure S1).

**Figure 2.**
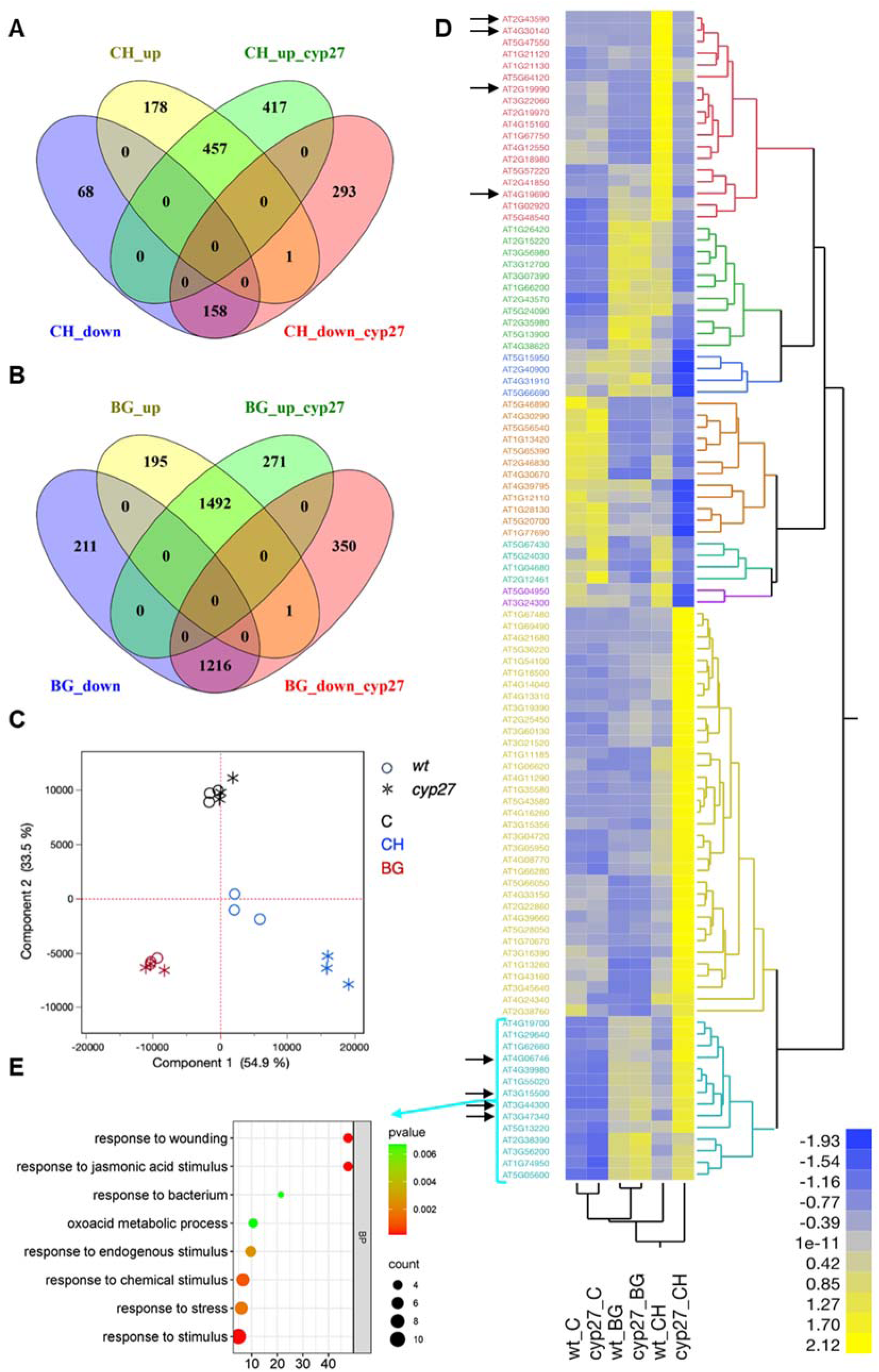
Effect of loss of *CYP71A27* on the transcriptome response of Arabidopsis roots to bacteria. **A** Venn diagram comparing the number of DEGs regulated by CH267 vs. mock in *cyp71A27* mutant and WT **B** Venn diagram comparing the number of DEGs regulated by *B. glumae* vs. mock in *cyp71A27* mutant and WT **C** PCA analysis of the transcriptome of WT and *cyp71A27* (cyp27) roots treated with mock (C), *Pseudomonas* sp. CH267 (CH) and *Burkholderia glumae* PG1 (BG). **D** Heat map of the mean expression of DEGs of the interaction effect of cyp27 and CH267. Genes selected as candidates for further analyses are marked with arrows. **E** GO enrichment analysis of DEGs corresponding to cluster 8 of the interaction cyp27 with CH267.

Interestingly, among the genes induced by *B. glumae* in both genotypes we detected several marker genes for sulfur deficiency, the *RESPONSE TO LOW SULFUR LSU1* and *LSU3*, *SERINE HYDROXYMETHYLTRANSFERASE SHM7*, *GAMMA-GLUTAMYL CYCLOTRANSFERASE GGCT2;1*, and *BETA GLUCOSIDASE BGLU28* (Ristova and Kopriva, 2022). Enrichment analysis using Virtual Plant (Katari et al., 2010) revealed that many Gene Ontology (GO) functional categories and KEGG metabolic pathways were shared among DEGs of the two bacterial treatments and/or both genotypes. Within the upregulated genes, functional categories associated with “response to other organism”, “indole-containing compound metabolic process” and “peroxidase activity” were enriched in all four datasets, while “suberin biosynthetic process”, or “glutathione transferase activity”, were enriched by both bacterial strains in WT but only by *B. glumae* in *cyp71A27* (Supplemental Figure S2; Supplemental Table S2). Similarly, both strains caused downregulation of genes assigned to categories “cell wall organization or biogenesis”, “central vacuole”, and “xyloglucan metabolic process” in both genotypes, while “photosynthesis”, and “water channel activity” only in the WT (Supplemental Figure S2).

Interestingly, several GO terms related to isoprenoid metabolism were enriched among genes downregulated in mock treated *cyp71A27* compared to WT, in particular “thalianol metabolic process” (Supplemental Table S2). The GO term analysis was confirmed by finding similar KEGG pathways enriched among the DEGs (Supplemental Table S3). Since the two bacteria trigger different effects in the plant, we were interested whether we find genes regulated in an opposite manner and whether they might help to explain the different plant responses. Indeed, in the WT 37 genes were downregulated by *B. glumae* and upregulated by CH267, including 4 genes for peroxidases, sugar transporter *STP4*, and nitrate transporter *NRT1;5*. On the other hand, 7 genes were downregulated in CH267 treated roots and induced by the pathogen, for example, *Ferredoxin 1*, *Chalcone synthase TT4* and *Flavonol synthase FLS1* (Supplemental Figure S1; Supplemental Table S1). In *cyp71A27* 60 DEGs were upregulated by CH267 and downregulated by *B. glumae* and 23 DEGs were *vice versa* repressed by CH267 and induced by *B. glumae* (Supplemental Figure S1). Correspondingly, the “flavonoid biosynthetic process” was highly enriched among genes upregulated in *B. glumae* and downregulated in the PGP strain, represented by the *TT4* and *FLS1* genes (Table 1). On the other hand, “peroxidase activity” and “heme binding” are two GO terms overrepresented in DEGs upregulated in the CH267 and repressed in *B. glumae* (Table 1).

**Table 1.** Enrichment analyses of GO terms in DEGs from Arabidopsis roots upregulated in *B. glumae* and downregulated after treatment with the PGP strain *Pseudomonas* sp. CH267.

| GO term | Observed frequency | Expected frequency | Enrichment | adj. P value | Category | Genes |
| --- | --- | --- | --- | --- | --- | --- |
| response to oxidative stress | 5/34 14.7% | 247/22304 1.1% | 13.4 | 0.00490 | BF | AT5G19890 AT5G13930 AT1G49570 AT5G64100 AT2G18980 |
| lipid transport | 3/34 8.8% | 137/22304 0.6% | 14.7 | 0.03862 | BF | AT4G12550 AT3G53980 AT4G33550 |
| lipid localization | 3/34 8.8% | 153/22304 0.6% | 14.7 | 0.03862 | BF | AT4G12550 AT3G53980 AT4G33550 |
| flavonoid biosynthetic process | 2/34 5.8% | 46/22304 0.2% | 29.0 | 0.04573 | BF | AT5G13930 AT5G08640 |
| flavonoid metabolic process | 2/34 5.8% | 51/22304 0.2% | 29.0 | 0.04841 | BF | AT5G13930 AT5G08640 |
| cell wall | 5/27 18.5% | 458/19822 2.3% | 8.0 | 0.00729 | CC | AT5G07030 AT5G47550 AT5G38940 AT5G64100 AT5G64120 |
| external encapsulating structure | 5/27 18.5% | 462/19822 2.3% | 8.0 | 0.00729 | CC | AT5G07030 AT5G47550 AT5G38940 AT5G64100 AT5G64120 |
| oxidoreductase activity, acting on peroxide as acceptor | 5/41 12.1% | 103/24443 0.4% | 30.3 | 0.00004 | MF | AT5G19890 AT1G49570 AT5G64100 AT5G64120 AT2G18980 |
| peroxidase activity | 5/41 12.1% | 103/24443 0.4% | 30.3 | 0.00004 | MF | AT5G19890 AT1G49570 AT5G64100 AT5G64120 AT2G18980 |
| antioxidant activity | 5/41 12.1% | 130/24443 0.5% | 24.2 | 0.00008 | MF | AT5G19890 AT1G49570 AT5G64100 AT5G64120 AT2G18980 |
| electron carrier activity | 5/41 12.1% | 427/24443 1.7% | 7.1 | 0.01756 | MF | AT5G19890 AT1G49570 AT5G64100 AT2G18980 AT1G10960 |
| heme binding | 4/41 9.7% | 290/24443 1.1% | 8.8 | 0.02632 | MF | AT5G19890 AT1G49570 AT5G64100 AT2G18980 |
| tetrapyrrole binding | 4/41 9.7% | 314/24443 1.2% | 8.1 | 0.02632 | MF | AT5G19890 AT1G49570 AT5G64100 AT2G18980 |
| iron ion binding | 4/41 9.7% | 352/24443 1.4% | 6.9 | 0.03354 | MF | AT5G19890 AT1G49570 AT5G64100 AT2G18980 |
| lipid binding | 3/41 7.3% | 173/24443 0.7% | 10.4 | 0.03354 | MF | AT4G12550 AT3G53980 AT4G33550 |

To better understand the specific transcriptional changes in the *cyp71A27* mutant under CH267 treatment we determined the main effect of the loss of *CYP71A27* and its interaction with treatments with *B. glumae* and CH267 using DESeq2 (Figure 2D, Supplemental Figure S3, Supplemental Table S1). This analysis identified 66 genes as the main effect of the *cyp71A27* genotype and 9 and 100 DEGs in the interaction effects with *B. glumae* and CH267, respectively (Supplemental Figure S3). The most interesting were found to respond in the interaction between mutant and CH267 and can be divided into 8 clusters according to their expression profile (Figure 1D). The genes less expressed in CH267 treated *cyp71A27* than WT form 6 clusters, without a clear functional enrichment. The effect of *cyp71A27* mutation is the strongest in cluster 1, which contains genes, such as iron transporter *IRT1* (AT4G19690), pathogenesis-related protein *PR1-LIKE* (AT2G19990), chitinase family protein (AT2G43590), and GDSL-like Lipase/Acylhydrolase superfamily protein *CDEF1* (AT4G30140). On the other hand, clusters 7 and 8 contain genes that are higher expressed in the mutant compared to WT after CH267 treatment and show a clear functional enrichment of GO terms for metabolism of organic compounds, like “oxoacid metabolic process”, or jasmonate related categories (Figure 2D, E). Cluster 8 includes genes for two transcription factors, *ANAC055* (AT3G15500) and *RAP2.9* (AT4G06746), a regulator of jasmonate signalling *JAZ10* (AT5G13220), the lipoxygenase *LOX1* (AT1G55020), nitrilase *NIT2* (AT3G44300), and asparagine synthase *ASN1* (AT3G47340) (Figure 2E). Altogether, the transcriptional analysis pointed to a specific function of CYP71A27 in the interaction with CH267 and suggested an involvement of jasmonate controlled processes in this interaction. Since both bacterial strains induce the biosynthesis of camalexin we specifically explored the regulation of the corresponding genes. Both bacterial strains highly induced the gene for the key enzyme in camalexin synthesis *CYP71A12*. Other genes for camalexin synthesis were strongly upregulated by *B. glumae* while CH267 induced these genes as well, but to a lower extent (Supplemental Figure S4). The extent of this regulation thus corresponds to the accumulation of camalexin, which is upregulated in Arabidopsis by treatment with both bacteria but to a much greater extent with *B. glumae* irrespective of plant genotype (Supplemental Figure S4).

### Novel genes for plant interaction with PGP bacteria

Given the loss of growth promotion by CH267 in *cyp71A27* mutant (Koprivova et al., 2019) and the specific differential regulation of genes in clusters 1 and 8 only by CH267 (Figure 2D), we considered these genes as the prime candidates for the downstream effect(s) of the loss of CYP71A27 and selected several ones for detailed analysis, namely *Chitinase* AT2G43590, GDSL-like Lipase *CDEF1*, *PR1-LIKE*, and *IRT1* in cluster 1 as well as *ANAC055*, *RAP2.9*, *NIT2*, and *ASN1* in cluster 8. We first verified their expression in the WT and *cyp71A27* roots after treatments with CH267, *B. glumae*, and mock by qRT-PCR. As expected from the RNAseq analysis, all four genes from cluster 1 were less upregulated by CH267 in the mutant than in WT, while those from cluster 8 showed the opposite pattern (Supplemental Figure S5). We obtained homozygous T-DNA lines for these eight genes and incubated them with the CH267. As the primary readout we determined camalexin in the shoots and compared with the *cyp71A27* mutant. Indeed, after treatment with CH267, camalexin concentration was significantly lower in *cyp71A27* mutant, compared to WT, but interestingly, the same was true for three of the cluster 1 mutants and all four mutants from cluster 8, the exception being the *cdef1* (Figure 3A). Thus, mutants in the genes from both cluster 1 and cluster 8 seem to phenocopy *cyp71A27* in the camalexin response to treatment with the PGP strain CH267.

**Figure 3.**
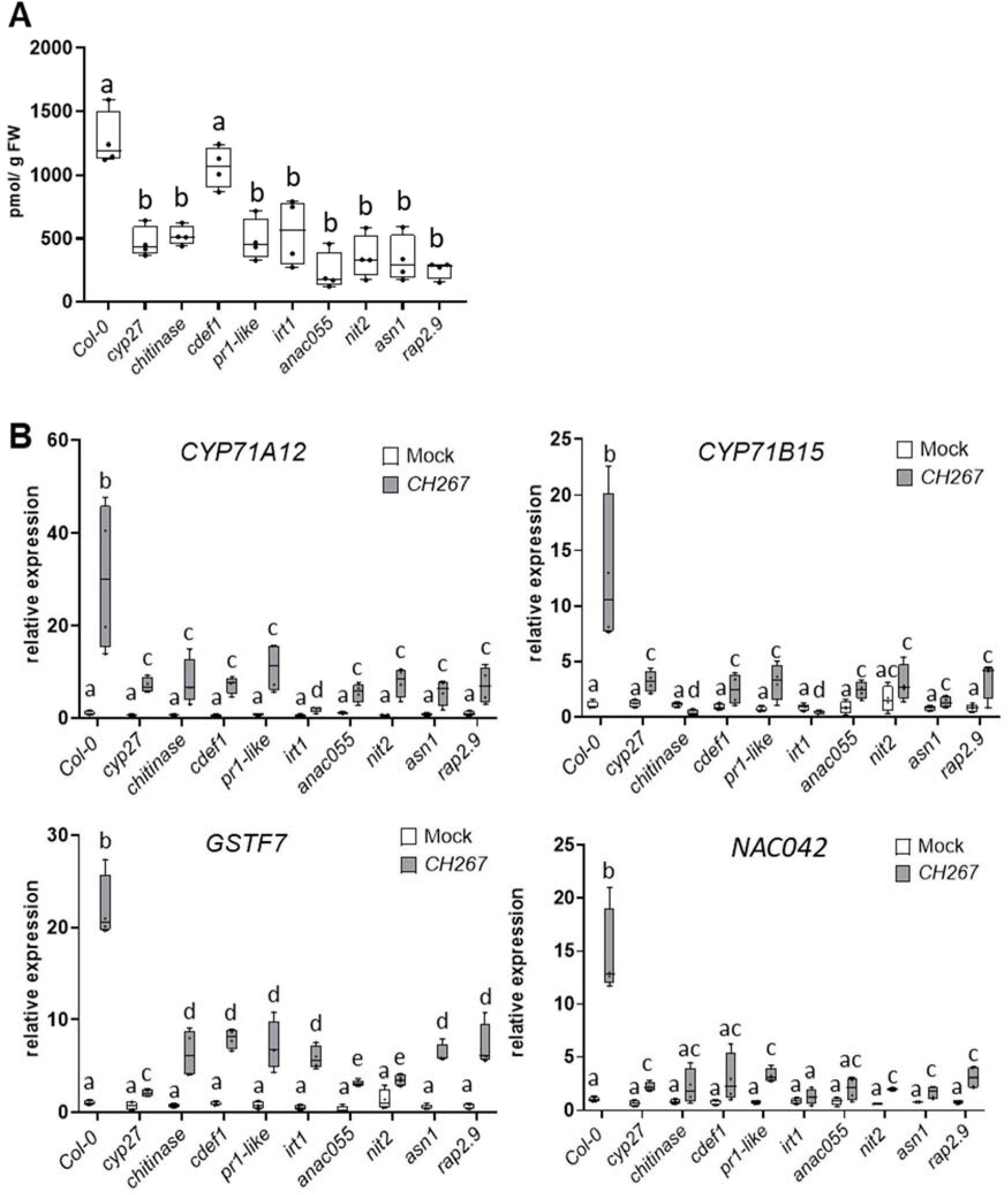
Analysis of mutants in candidate genes for response to CH267. **A** Camalexin levels in shoots of WT, *cyp71A27*, and mutants in cluster 1 and cluster 8 genes connected to *CYP71A27* – *chitinase, cdef1, pr1-like, irt1, anac055, nit2, asn1,* and *rap2.9* treated with CH267. **B** Relative transcript levels of marker genes responding to bacterial infection - *CYP71A12, CYP71B15, GSTF7, NAC042* – in roots of WT, *cyp71A27*, and mutants in cluster 1 and cluster 8 genes treated with mock or CH267. The qRT-PCR results were obtained from 4 biological and 2 technical replicates, the transcript levels of WT Col-0 in mock treated roots was set to 1. Letters mark values significantly different at P<0.05 (ANOVA).

In the next step we performed a whole-transcriptome analysis on roots of the mutants treated with the CH267 and mock and compared expression of several marker genes for response to bacteria. The key gene in camalexin synthesis pathway, *CYP71A12*, was highly induced in the roots of WT, and this induction was much lower in *cyp71A27* (Figure 3B). After treatment with CH267, all eight mutants showed a slight but significant upregulation of *CYP71A12*, similar to *cyp71A27*. The same pattern was shown for *CYP71B15* (*PAD3*), another gene in camalexin synthesis, highly induced by CH267 in the WT but less responsive in *cyp71A27*, while all eight mutants in the candidate genes showed a dampened induction when compared to WT (Figure 3B). Interestingly, in *irt1* the *CYP71B15* was actually downregulated by CH267, and *CYP71A12* was even less responsive that in *cyp71A27*. Regulation of another two marker genes, *GSTF7* and *NAC042*, which controls camalexin synthesis (Saga et al., 2012), showed the same pattern as *CYP71A12* and *CYP71B15,* with the genes being upregulated to much lower extent in *cyp71A27* than in the WT, and the mutants showing similar low upregulation as the *cyp71A27* (Figure 3B). Thus, the mutants in the 8 candidate genes from a large part phenocopy *cyp71A27* also in the alterations in transcriptional response to CH267, and seem, therefore, to be part of the same regulatory network. Interestingly, this similarity is conserved in mutants of genes from both clusters 1 and 8 despite their opposite expression pattern in *cyp71A27* and WT.

### Response of proteomes of cyp71A27 and WT Arabidopsis roots to pathogenic and PGP bacterial strains

Next we asked how far the changes we detected in transcripts are detectable on the proteome level. We analysed the root and shoot proteomes of WT Col-0 and *cyp71A27* mutant in all three conditions, mock and treatments with CH267 and *B. glumae*. In the WT, as suggested by the results of RNAseq (Figure 2), and using stringent criteria, more differentially abundant proteins (DAP; q<0.01 and FC>2) were found after treatment with *B. glumae* than with CH267, both in roots and shoots (Figure 4A). Analogously to the transcriptome analysis, while some proteins were affected by both bacteria, more DAPs were specific for one strain, especially in the case of *B. glumae* (Figure 4A). Similarly, in the roots of *cyp71A27* about half of the DAPs more abundant after treatment with CH267 was also more abundant after the pathogen strain, while in the shoots the numbers of shared and specific DAPs were similar for both bacteria (Supplemental Figure S6A). PCA revealed that in the root proteome, treatment with *B. glumae* showed more substantial effects on both WT and *cyp71A27* (Figure 4B), as before on the transcriptome level. Interestingly, the shoot proteomes showed the effects of the bacterial treatments compared with mock, however, the effects of the bacterial strains were not clearly separated. On the other hand, although *CYP71A27* is primarily expressed in the roots (Koprivova et al., 2019), the shoot proteomes of the mutant and WT were clearly separated within each treatment (Figure 4B). Comparing proteomes of Col-0 and the *cyp71A27* mutant under all three conditions showed that in the shoots, more DAPs with increased abundance than lower abundance were found in the treated plants compared to mock, while in the roots the higher and lower abundant proteins were more in balance (Figure 4C). In roots of the *cyp71A27* mutant the number of DAPs were greatly reduced compared to Col-0, while the difference was much lower in the shoots (Figure 4C).

**Figure 4.**
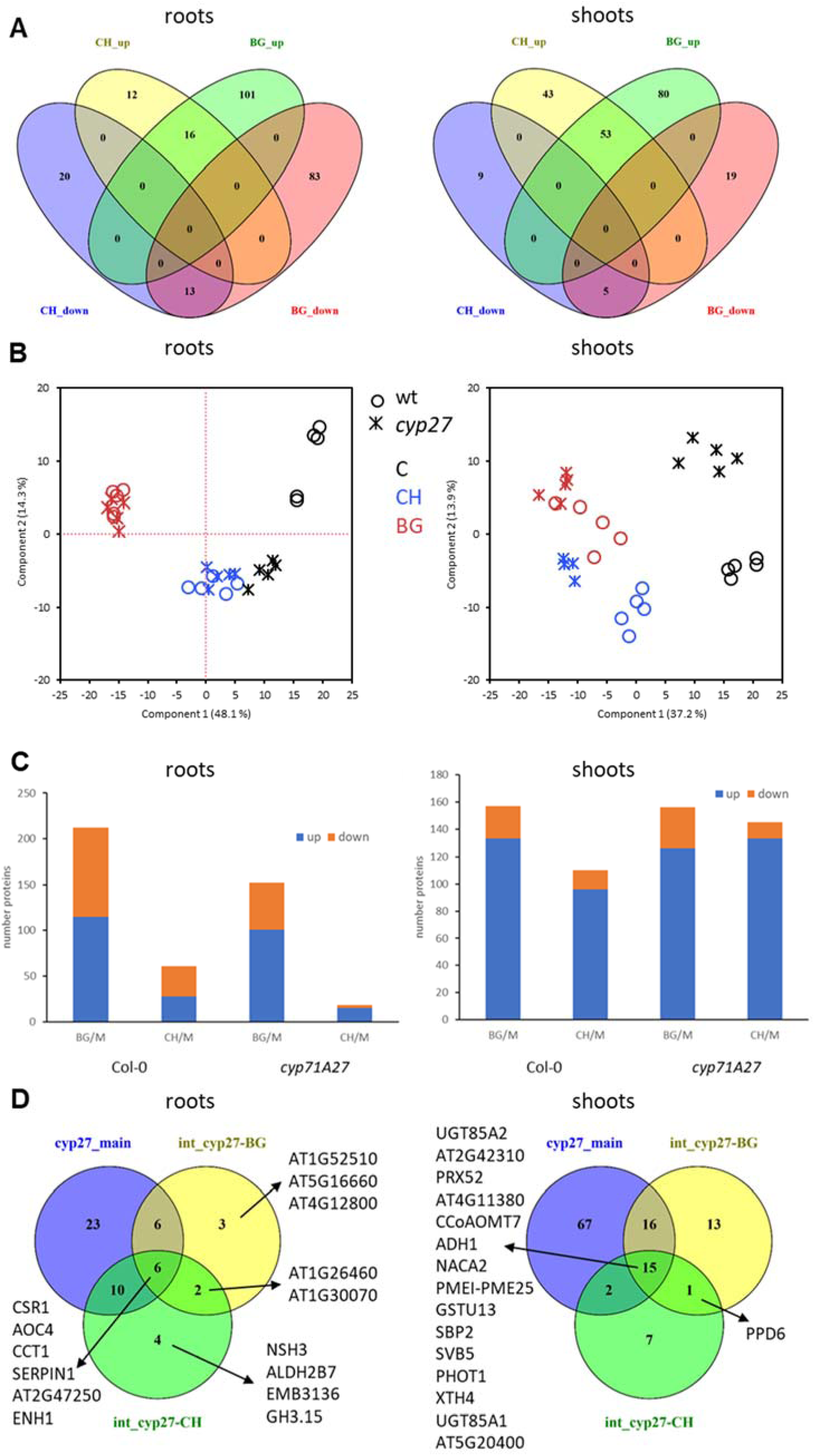
Proteome response of Arabidopsis to *Pseudomonas* sp. CH267 (CH) and *Burkholderia glumae* PG1 (BG). **A** Venn diagrams showing the number of DAPs in WT Col-0 roots and shoots after treatments with CH and BG. **B** PCA analysis of the roots and shoots proteomes of WT Col-0 and *cyp71A27* (cyp27) roots treated with mock (C), *Pseudomonas* sp. CH267 (CH) and *Burkholderia glumae* PG1 (BG). **C** Number of upregulated and downregulated DAPs in *cyp71A27* and WT roots and shoots. **D** Venn diagrams showing the number of DAPs in *cyp71A27* main effect, and interaction of cyp27 with CH and BG.

The response to *B. glumae* was largely similar in the roots of *cyp71A27* and the WT (Supplemental Figure S6B). Enrichment analysis revealed that *B. glumae* induced proteins involved in “response to biotic stimulus” but also number of proteins involved in metabolism, such as “aspartate family amino acid catabolic process”, “indole-3-acetonitrile nitrilase activity”, or “glutathione binding”. Among the proteins less abundant in *B. glumae* treated roots several categories connected with protein translation were enriched. The number of categories enriched in roots treated with CH267 was much lower, especially in *cyp71A27* (Supplemental Figure S7, Supplemental Table S5). The less abundant DAPs in the WT included processes connected to cytoskeleton, but also included proteins connected to chloroplasts, as observed in the RNAseq data (Supplemental Figure S7, Supplemental Table S5). Correspondingly, among the *B. glumae* induced DAPs the KEGG pathways “Amino Acid Metabolism”, “Valine, leucine and isoleucine degradation”, “Biosynthesis of Other Secondary Metabolites”, and “Phenylalanine metabolism” were enriched, the latter two also in the 16 proteins upregulated by both bacteria (Supplemental Table S6).

In the shoots, the bigger part of DAPs was common in *cyp71A27* and WT, whereas the less abundant DAPs were mostly genotype specific. Accordingly, in both genotypes the DAPs higher abundant by treatments with both strains were enriched in similar GO categories as in the roots (Supplemental Figure S7, Supplemental Table S5) and in the KEGG pathway “amino acid metabolism” (Supplemental Table S6). As suggested already by the enrichment of GO terms, among the CH267 induced DAPs the pathways of “Phenylalanine metabolism” and “Phenylpropanoid biosynthesis” were induced, while among the *B. glumae* induced DAPs it was the “Phenylalanine, tyrosine and tryptophan biosynthesis”, again similar to the roots (Supplemental Table S6). Given the role of camalexin in plant microbe interaction, it is worth to mention that CYP71A12 was not significantly changed in the roots but detectable in the shoots only after bacterial treatments, and significantly more abundant in *B. glumae* treated plants of both genotypes (Supplemental Figure S8).

In the WT and *cyp71A27*, 38 and 40 DAPs, respectively, were common in shoots and roots after treatment with *B. glumae* while only 6 and 4 proteins were common in both organs in plants treated with CH267 (Supplemental Figure S9). In the control, mock treated plants, the loss of *CYP71A27* resulted in differential abundance of 19 and 51 proteins in roots and shoots, respectively, none of which was found in both tissues (Supplemental Table S4). Then, as with the transcriptomics, we determined the main effect of the loss of *CYP71A27* as well as its interaction with the two bacterial treatments. In the roots, 6 proteins were regulated in all three conditions (Figure 4D). The DAPs belonging to the cyp27*CH267 interaction term were particularly enriched in the GO term “cytidylyltransferase activity” but also in “lipid biosynthetic process” and “plastid stroma” (Supplemental Table S5). The interaction term of *cyp71A27* with *B. glumae* on the other hand contained DAPs that belonged to various GO terms connected to plastids, such as “plastid envelope” and “thylakoid membrane”. In the shoots, the analysis revealed 100 DAPs as the main *cyp71A27* effect, and 25 and 45 DAPs in interaction terms with CH267 and *B. glumae*, respectively (Figure 4D). Fifteen DAPs were found in all three categories, such as the peroxidase PRX52, glutathione transferase GSTU13, or ABA responsive SVB5 protein.

We then compared the DEGs and DAPs in the roots. In the WT there was a high proportion of DAPs induced by *B. glumae* and CH267, that corresponded to DEGs upregulated in the RNAseq analysis, 50.4% and 42.8%, respectively (Supplemental Figure S10; Supplemental Table S7). On the other hand, the overlap between less abundant DAPs and DEGs was much smaller with 17.7% and 15.2% in *B. glumae* and CH267 treated roots, respectively (Supplemental Figure S10). In the roots of *cyp71A27* treated with CH267, 13 out of 15 more abundant DAPs and 1 of the 3 less abundant DAPs were regulated in the same way as the corresponding genes. Also the portion of DAPs regulated by *B. glumae* on transcriptional level was higher in *cyp71A27* than in WT (Supplemental Figure S10, Supplemental Table S7). The high level of overlap between DAPs and DEG is reflected also in the enrichment analyses, since, for example, GO terms, such as “glutathione binding”, “glutathione transferase activity”, and “indole-containing compound biosynthetic process” were found enriched among both upregulated DEGs and DAPs after treatment with *B. glumae* (Supplemental Table S5). Thus, the physiological changes triggered by the two bacterial strains are regulated on both transcriptional and posttranscriptional levels.

### Treatment with CH267 and B. glumae affects metabolite composition of the roots and exudates

In the next step we quantified metabolites from the roots and shoots of *cyp71A27* and WT treated with the two bacterial strains or mock. First we determined the concentration of several sulfur-containing metabolites, because they are related to camalexin synthesis and function and camalexin is induced by both bacteria in shoots and roots (Supplemental Figure S4B), and because of the upregulation of genes involved in S homeostasis by *B. glumae*. The glucosinolates are a class of secondary metabolites involved in immunity, like camalexin, and GO terms connected to glucosinolates were enriched among both DEGs and DAPs. The aliphatic, methionine derived glucosinolates were not affected by neither the bacterial treatment nor the loss of *CYP71A27* (Supplemental Figure S11A). On the other hand, after treatment with *B. glumae* the content of indolic glucosinolates, derived from tryptophan, was lower in the roots of the *cyp71A27* mutant than in WT (Figure 5A). In the WT, cysteine concentration doubled in roots treated with CH267 and tripled after incubation with *B. glumae*, but in shoots was increased only after pathogen treatment (Figure 5B, Supplementary Figure S11). Loss of CYP71A27 had only small impact in the roots, where after treatment with *B. glumae* Cys content was slightly lower than in WT, however, in shoots Cys increased in mock and CH267 treated mutants compared to WT (Supplemental Figure S11B). GSH concentration was increased in the roots of WT by both strains, more by *B. glumae* than by CH267, similar as Cys (Figure 5C). In *cyp71A27* roots GSH was induced less strongly by *B. glumae* and not at all by CH267. In the shoots, GSH was increased by *B. glumae* in both genotypes, but by CH267 only in the *cyp71A27*. Thus, both PGP and pathogen bacteria trigger alterations in concentrations of S-containing compounds and the CYP71A27 influences this response and affects also the coordination between roots and shoots.

**Figure 5.**
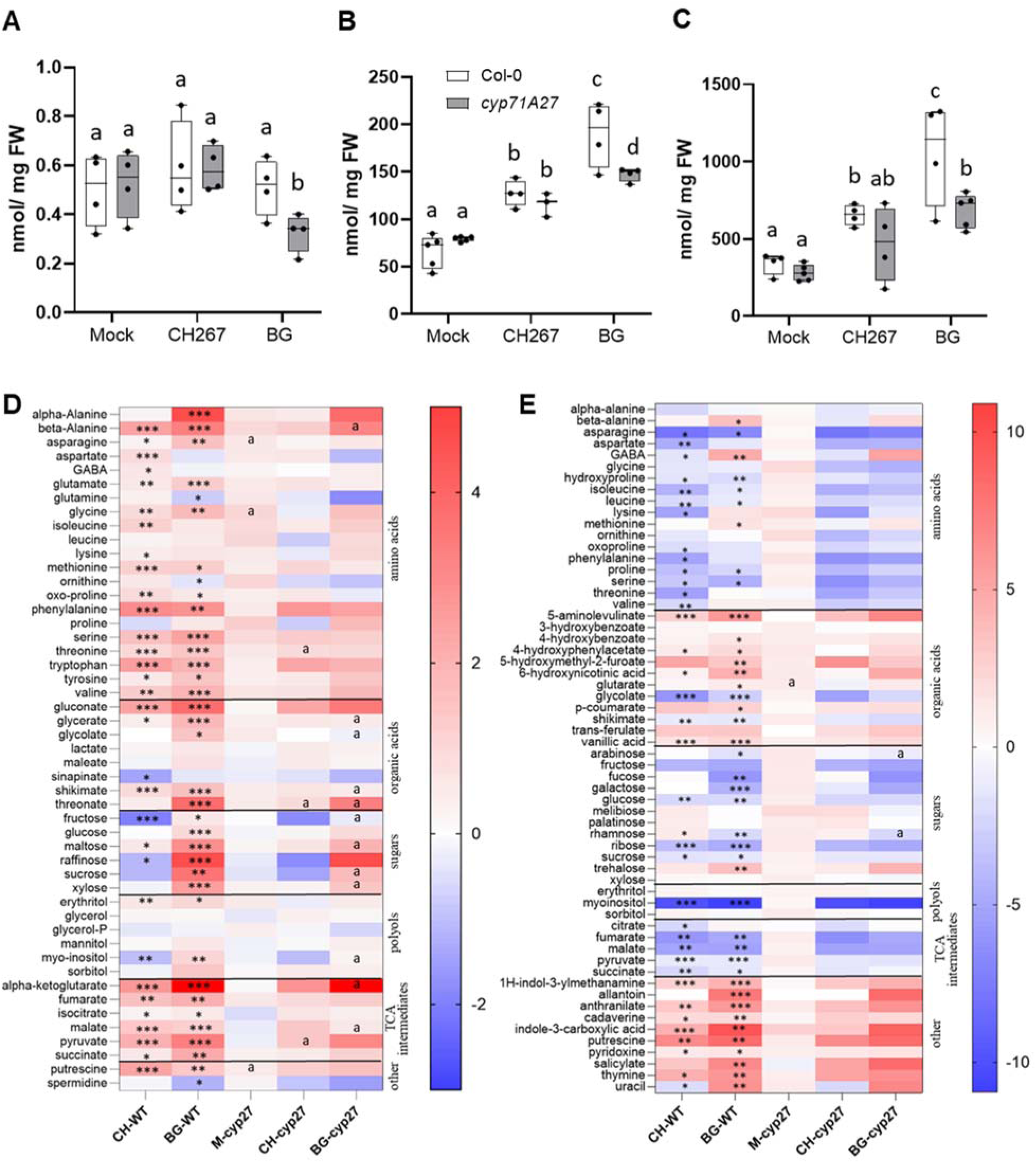
Metabolite analysis of response of Arabidopsis to *Pseudomonas* sp. CH267 (CH) and *Burkholderia glumae* PG1 (BG). The contents in roots of (**A**) indolic glucosinolates, (**B**) cysteine, and (**C**) glutathione were determined by HPLC. Different letters mark values dignificantly different at p<0.05 (T-test, n=4). **D** Metabolites were determined in WT and *cyp71A27* roots treated with mock, CH, or BG by GC-MS. **E** Exudates were collected from the treated plants and subjected to metabolite analysis by GC-MS. The heat maps show the log2 of fold change between bacteria treated plants and mock, as well as mock treated *cyp71A27* vs. WT (M-cyp27). Asterisks represent significant differences between bacterial treatments and mock in WT (T-test, * p<0.05, **p<0.01, *** p<0.001). Letter „a“ marks values different between *cyp71A27* and WT at p<0.01 (T-test, n=5).

To get a better insight into the effects of the bacterial treatments on plant metabolites, we further quantified 50 primary metabolites, mainly amino acids, carbohydrates, and organic acids in the roots. Most of these metabolites showed significant alterations in their abundance by at least one of the strains, again, with *B. glumae* having a stronger effect (Figure 5D). Also, most of the metabolites were induced by both strains with the exception of sugars, which were mostly more abundant after *B. glumae* treatment but not affected or downregulated by CH267 (Figure 5D). The highest induction was observed for roots treated with *B. glumae*, where, e.g., α-ketoglutarate was induced 154-fold in the WT and alanine more than 20-fold (Figure 5D, Supplemental Table S8). Among the metabolites induced by both strains to a similar level there are the precursors of camalexin and glucosinolates, i.e. tryptophan and methionine, corresponding to the results of transcriptomics and proteomics. The same pattern of regulation was shared by several other amino acids, i.e. phenylalanine, serine and threonine, and by TCA cycle intermediates fumarate, isocitrate, and malate (Figure 5D). The metabolites more strongly induced by *B. glumae* than by CH267 include besides α-ketoglutarate and alanine most prominently the amino acids ß-alanine, valine, asparagine, and tyrosine, as well as glycerate, gluconate, and the sugars glucose and xylose. Among the sugars, raffinose, sucrose, and fructose were significantly less accumulated in CH267 treated roots than in mock, while *B. glumae* induced these metabolites (Figure 5D, Supplemental Table S8).

The effect of loss of CYP71A27 on the root metabolites was not very prominent without the bacterial treatment. Only putrescine, asparagine, and glycine were slightly more abundant in the roots of mock treated *cyp71A27* compared to WT (Figure 5D). In the roots treated by CH267, threonate was only induced in *cyp71A27*, while threonine was induced in both genotypes but more in *cyp71A27* and pyruvate was induced less in the mutant. The *cyp71A27* differed more from WT after treatment with *B. glumae*, with many metabolites being induced to a lower extent than in the WT, including sucrose, maltose, xylose, α-ketoglutarate, shikimate, glycerate, and ß-alanine. The only metabolite affected differently in the mutant by both bacteria was threonate, which in the pathogen treated roots was induced less that in WT, opposite to treatment with CH267 (Figure 5D). Thus, unlike for transcriptome and proteome, the effect of the loss of *cyp71A27* on metabolome was larger after treatment with *B. glumae* than with CH267.

For the plant communication with microbes, the metabolites exuded from roots are of the utmost importance (Sasse et al., 2018). Therefore, we also analysed the metabolites in root exudates, extracted from the nutrient solution after 3 days of incubation with the two bacterial strains or mock. Sixty one metabolites were detected by the GC-MS method, about a half of them overlapping with the metabolites determined in the root. Most of the metabolites were exuded even without bacterial treatment, except trans-ferrulic acid, 5-aminolevulinic acid, and allantoin (Supplemental Table S8). Interestingly, while treatment with the bacteria led mostly to higher concentration of metabolites in the roots, in the case of exudates about half of the metabolites were less abundant, especially the amino acids, sugars, and TCA intermediates. The metabolites found to a much greater extend in the exudates after treatment with both strains include indole 3-carboxylic acid (ICA), putrescine, thymine, and anthranilate (Figure 5E). On the other hand, we found highly reduced concentrations of myo-inositol, asparagine, malate, and fumarate. Interestingly, two metabolites, γ-aminobutyrate and uracil, were less abundant in exudates of CH267 treated plants than in mock but increased after treatment with *B. glumae*. The loss of CYP71A27 had almost no effect on the exudation pattern, only glutarate was exuded in greater amount from mock treated *cyp71A27* while arabinose and rhamnose were more strongly repressed by *B. glumae* in WT than in *cyp71A27* (Figure 5E; Supplemental Table S8). Using a targeted analysis, camalexin was found only in exudates from bacteria treated roots, more after incubation with *B. glumae*, as reported previously (Koprivova et al., 2023) and surprisingly, its levels were not different in *cyp71A27* compared to WT (Supplemental Figure S12). Overall, while both bacteria strongly affected metabolite concentrations in the roots and exudates, the effects of the loss of CYP71A27 were less clear on the metabolite level than on transcriptome and proteome.

### Effect of treatment with CH267 and B. glumae on Arabidopsis ionome

Since the bacterial treatment induced several genes related to nutrient uptake and homeostasis, such as *IRT1*, Copper transport protein family (*AT5G52710*), phosphate transporter *PT3;2*, or *NRT1;8* (Supplemental Table S1), we assessed whether the incubation of plants with the bacterial strains affects nutrient contents in the shoots. Indeed, with exception of sodium, all analysed elements were affected either by the bacterial treatment in the WT or accumulated differently in the *cyp71A27* mutant and WT (Figure 6). Boron and magnesium were less abundant after treatment with both strains, while potassium was higher in CH267 treated plants and lower in *B. glumae* WT plants compared to mock. Iron, sulfur, and zinc accumulated to higher levels in CH267 treated plants, while manganese after inoculation with *B. glumae* (Figure 6). The loss of CYP71A27 affected specifically phosphorus, calcium, and molybdenum content; these elements were not affected by the bacterial treatments in WT but P and Mo were lower in mock and *B. glumae* treated *cyp71A27* plants than in WT, while Ca was increased in the mutant irrespective of the treatment. Interestingly, the increased accumulation of Fe after CH267 treatment was lower in the mutant than in WT (Figure 6), corresponding to the regulation of *IRT1* (Figure 2D, Supplementary Figure S5).

**Figure 6.**
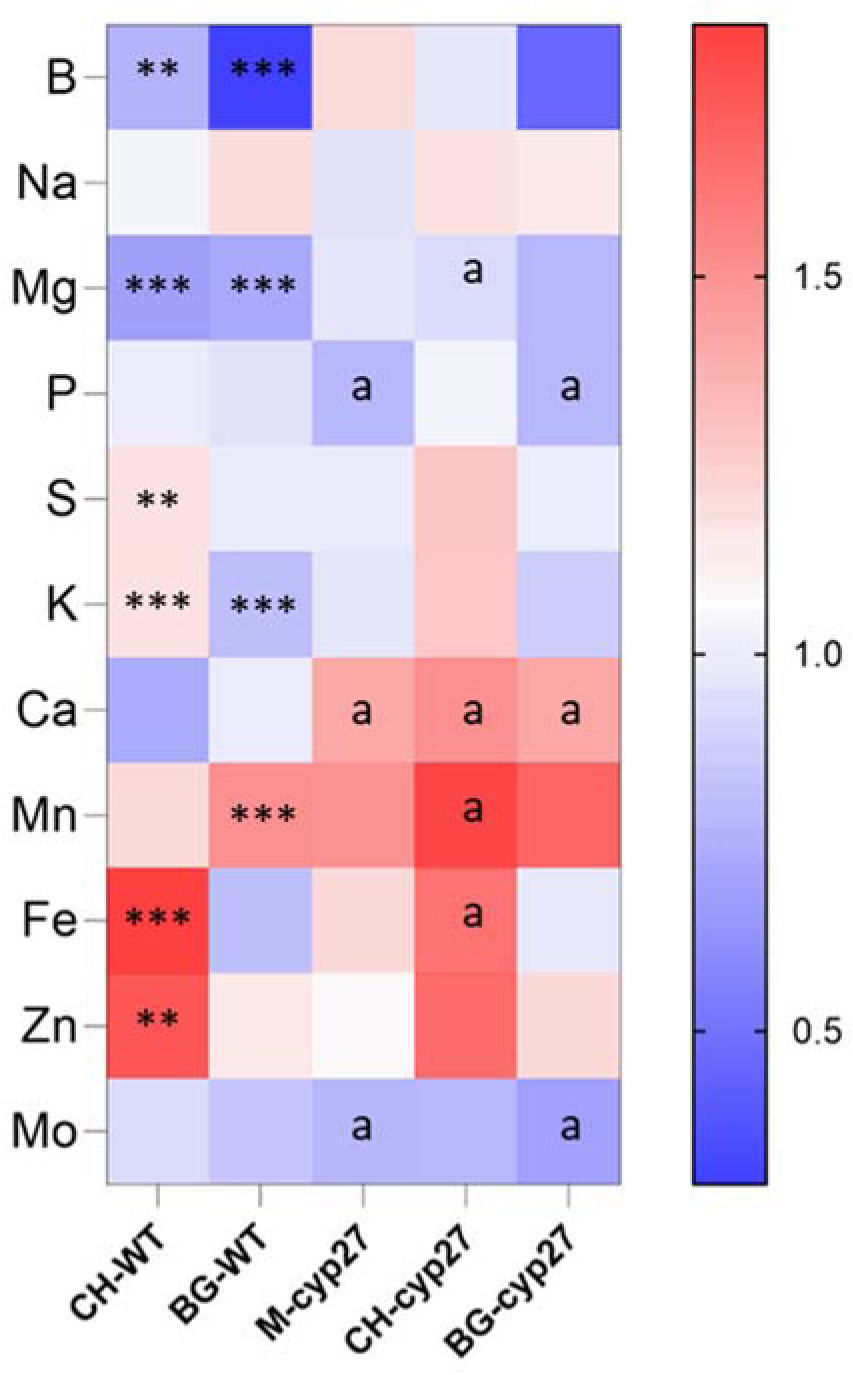
Response of Arabidopsis ionome to *Pseudomonas* sp. CH267 (CH) and *Burkholderia glumae* PG1 (BG). The contents of mineral elements were determined in shoots by ICP-MS. The heat map shows the log2 of fold change between bacteria treated plants and mock, as well as mock treated *cyp71A27* vs. WT (M-cyp27). Asterisks represent significant differences between bacterial treatments and mock in WT (T-test, ** p<0.01, *** p<0.001). Letter „a“ marks values different between *cyp71A27* and WT at p<0.01 (T-test, n=6).

### Network analysis

To assess to which extent the changes in gene expression in *cyp71A27* correlate with changes in metabolite levels we constructed interaction networks from the three comparisons, the *cyp71A27* main effect and the interactions *cyp71A27* with CH267 and with *B. glumae*. The correlation analyses resulted in distinct networks of different sizes for the three conditions (Supplemental Figure S13). The network with the fewest correlations was found for the *cyp71A27*B. glumae* interaction, while the interaction with CH267 produced the largest network. Interestingly, the networks showed a pronounced connectivity of the metabolites (Supplemental Figure S13), therefore, to focus on the most important correlations we limited the number of metabolites for the network construction to the 8 most highly regulated ones: camalexin, glutathione, cysteine, asparagine, pyruvate, sucrose, fructose, and myo-inositol. These subnetworks confirmed strong interconnection of the metabolites and only limited links to the genes (Figure 7). Thus, the only gene from the interaction between *cyp71A27* and *B. glumae* correlating with the 8 metabolites was AT3G21770, encoding a peroxidase PER30. The main effect of the loss of CYP71A27 highlights the interconnection of the metabolites, with central roles for camalexin, glutathione, and pyruvate, but also direct correlation of several genes (Figure 7A). Two genes are positively correlated with these three metabolites, AT2G37280 and AT5G38000, encoding the ABC transporter PDR5 and a Zinc-binding dehydrogenase, respectively. Other highly connected genes are AT2G40080 for EARLY FLOWERING 4 (ELF4), a component of a circadian clock machinery and AT5G48010 for thalianol synthase 1 (THAS1) (Figure 7A, Supplemental Table S9). The highest effect of the loss of CYP71A27 was observed after treatment with CH267, correspondingly, this network included the highest number of genes (Figure 7B). Also in this network, camalexin, GSH, and pyruvate were the most connected metabolites, as well as asparagine. The major hub genes were AT5G13900 and AT2G35980, for Bifunctional inhibitor/lipid-transfer protein/seed storage 2S albumin LPTG30 and YELLOW-LEAF-SPECIFIC GENE 9 (YSL9), respectively. Two genes for transcription factors connected to this network, the BHLH39 (AT3G56980) and MYB4 (AT4G38620), as well as genes for Glutamine Synthase GLN1;2, Chitinase, and Plant basic secretory protein (BSP) (Figure 7B). Altogether the network analysis dentified a number of new genes that might be connected to this response, like the genes coexpressed with *CYP71A27* described earlier.

**Figure 7.**
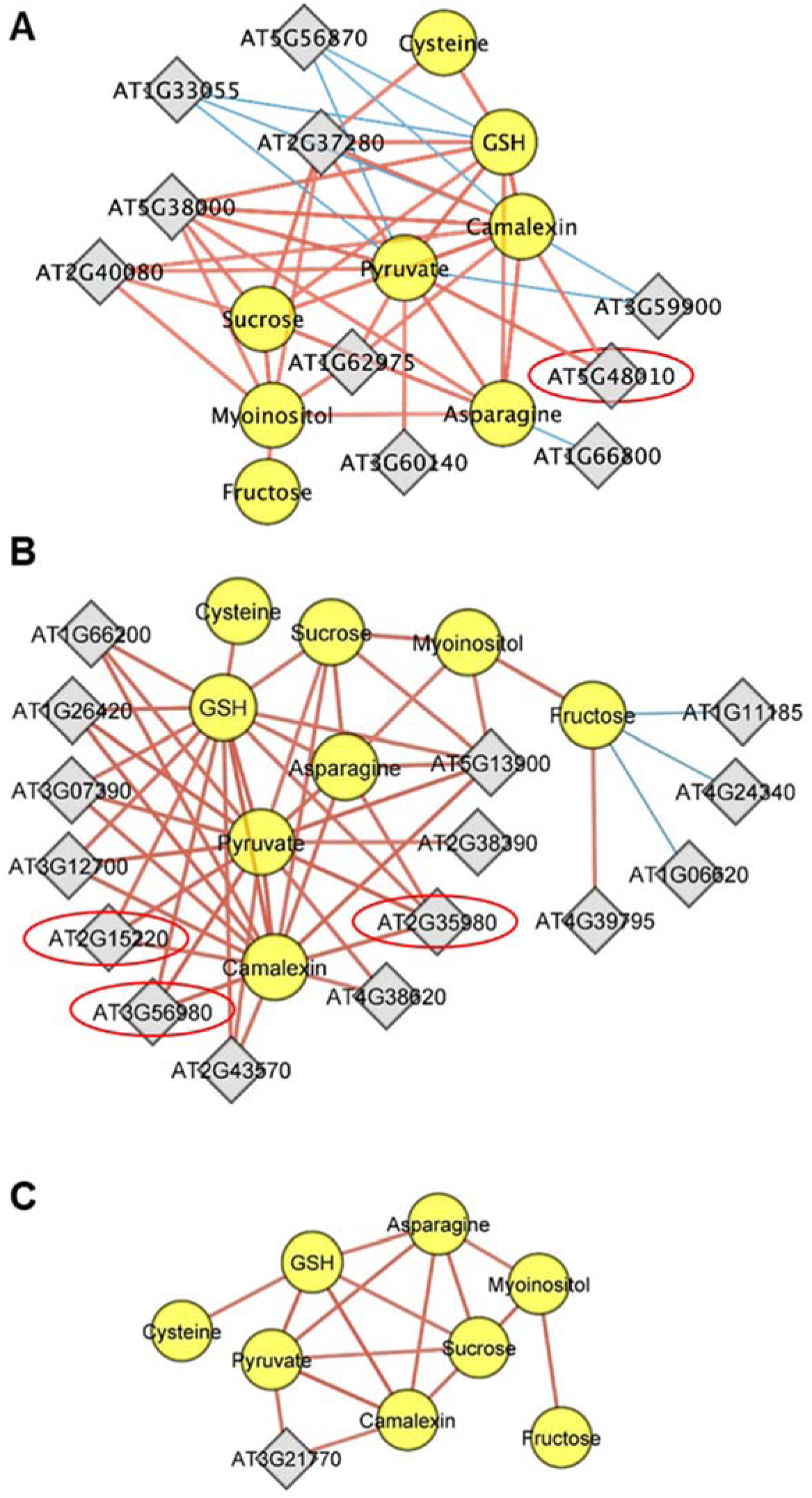
Network analysis of DEGs and 8 key metabolites. The networks were created by pairwise correlations between DEGs and metabolite levels and visualized in Cytoscape. Shown are subnetworks of 8 key metabolites responding most to the bacterial treatments from (**A**) *cyp71A27* main effect (**B**) interaction cyp27 and CH267, and (**C**) interaction cyp27 and *B. glumae* The genes are represented in gray rectangles and metabolites in yellow circles, brown lines represent positive correlation, and blue lines denote negative correlation. Full networks are shown in Supplemental Figure S14

Four of such genes were, therefore, analysed to more detail, namely *THAS1*, *YSL9*, *BSP*, and *BHLH39*. Initial analysis showed that while all four genes were induced in the roots by both bacterial strains, the upregulation by CH267 was much lower in the *cyp71A27* than in the WT, but *B. glumae* produced the same response in both genotypes (Supplemental Figure S5). Homozygous mutants were generated from T-DNA lines obtained from NASC and the mutants were analysed after treatment with CH267. Similar to the mutants in candidate genes from clusters 1 and 8, in the shoots of the four mutants camalexin accumulated to the same level as in *cyp71A27* and much less than in the WT (Figure 8A). Expression analysis of the mutants revealed that these mutants resemble *cyp71A27* also in the transcriptional response to CH267, since the four marker genes were less upregulated than the WT, with the exception of the two camalexin synthesis genes *CYP71A*12and *CYP71B15* in *bsp* (Figure 8B). Interestingly, in *thas1* and *bhlh39* two of the marker genes, *CYP71B15* and *NAC042*, were not induced by CH267 at all. Thus, also the mutants in genes from the *cyp71A27* network largely phenocopy the mutant in its low response to CH267. The network and the cluster analysis thus provided new genes with functions in the interaction of plants with plant growth promoting bacteria.

## DISCUSSION

### Multilevel response of A. thaliana to pathogen and PGP bacterial strains

Despite a great progress in deciphering the mechanisms of plant responses to individual microbes and microbial communities, the full understanding of how plants distinguish the commensal and pathogenic strains is not completely resolved (Ji et al., 2022; Paries and Gutjahr, 2023). In particular, while the roles of immunity are well established as crucial in plant reactions to different microorganisms (Teixeira et al., 2021), new components are still being identified (Koprivova et al., 2019). Studying how plant react to different microbiota using global omics technologies and the integration of the data on different levels has a great potential to identify the regulatory networks and new components of plant response to microbes (Jain et al., 2024). Therefore, we determined how Arabidopsis responds to the two contrasting bacterial strains on the multiple levels: transcriptome, proteome, and metabolome.

Numerous transcriptomics studies showed a large and differential effect of various bacterial strains on plant mRNA levels both in leaves and roots (Teixeira et al., 2021; Nobori et al., 2022). Even among commensal strains the variation is large and correspondingly small is the overlap forming a conserved response. Therefore, it was expected that the pathogen *B. glumae* affected expression of many more genes than the PGP strain (Figure 2). As with many other bacterial strains, treatments with both CH267 and *B. glumae* induced expression of genes connected to response to other organisms and, in fact, similar GO terms have been upregulated by the two bacteria as in roots treated with the flg22 peptide from bacterial flagellin (Teixeira et al., 2021). The overlap of the response was relatively high, only few functional categories were different, for example, categories related to ethylene and jasmonate signalling were induced by CH267, while those connected to abiotic stress and ABA were induced by *B. glumae* (Supplemental Table S2). Correspondingly, jasmonate and ethylene signalling were also induced during *Pseudomonas simiae* WCS417r induced systemic resistance against herbivores while tolerance against salt stress caused by *Enterobacter* sp. SA187 was dependent on ethylene (Pangesti et al., 2016; de Zelicourt et al., 2018). Also the upregulation of genes for suberin biosynthesis by CH267 is intriguing because alterations in the endodermal barrier system have a great impact on the plant root microbiome composition (Salas-Gonzalez et al., 2021). The 44 genes regulated in an opposite manner and hypothesised to explain the different response to the two strains, were enriched for genes involved in phenylpropanoid synthesis; in particular, flavonoid synthesis was upregulated after treatment with *B. glumae* and repressed by CH267 (Table 1).

Flavonoids are commonly found in root exudates and play important roles in plant interactions with other organisms (Ghitti et al., 2022). Remarkably, while there is only very small overlap in functional categories enriched in DEGs after treatment with *B. glumae* of Arabidopsis roots and rice leaves, the two genes involved in flavonoid metabolism *TT4* and *FLS1* were induced in both (Table 1; (Magbanua et al., 2014). Since flavonoids inhibit the production of the bacterial toxin toxoflavin in *B. glumae* (Castellanos et al., 2020), upregulation of flavonoid synthesis might be a specific mechanism of plant defence against *B. glumae*.

Apart of transcriptome changes, we were able to show that the two bacterial strains have a significant effect on plant proteome (Figure 4). This is important, because much less is known about plant response to bacteria on the protein level. The functional categories associated with the response to CH267 and *B. glumae*, particularly those related to defense and amino acid metabolism, correspond well to other studies. For example, incubation of Arabidopsis with PGPR strain *Paenibacillus polymyxa* E681 led to changes in abundance of proteins related to amino acid metabolism, antioxidant production, defense and stress responses (Kwon et al., 2016). Interestingly, after treatment with CH267 the enriched functional categories were conserved, but at the level of individual proteins the responses were different. The pathogenic strain, *B. glumae*, triggered more profound changes in the root proteome than the PGP strains, either CH267 in our study or other strains in the literature (Kwon et al., 2016; Witzel et al., 2017). Surprisingly, while we observed a significant overlap in DAPs between the treatment with *B. glumae* and CH267 in our study (Figure 4A), there were no overlapping proteins after treatment with a pathogenic protozoan *Plasmodiophora brassicae* despite both organisms primarily inducing changes in proteins related to defense (Devos et al., 2006). Remarkably, while in our study Arabidopsis roots were inoculated by the bacteria, there were significant changes in leaf proteomes (Figure 4).

This agrees with previous observations of induced systemic resistance, where rhizosphere bacteria protect plants also against leaf pathogens (Pieterse et al., 1996; Haney et al., 2018) and with significant changes in leaf proteome after inoculation with PGP strain WCS417r (Marzorati et al., 2023). Interestingly, even though the two PGP strains WCS417r and CH267 belong to the same genus, *Pseudomonas*, the changes they trigger in the leaf proteomes are different, with only two proteins, the glutathione transferases GSTF2 and GSTF7, being induced in leaves by both strains. This is unexpected, because comparison of treatments with CH267 and the pathogen *B. glumae* resulted in an overlap of 53 more abundant and 5 less abundant proteins (Figure 4). Thus, it seems that growth conditions have a major impact on the outcomes of the plant microbe interactions, at least in terms of differentially abundant proteins. It has to be noted, however, that we used very stringent conditions for the proteomics analysis, because we were interested in the most significant alterations. The overlaps might, therefore, be larger under the usual less stringent fold change threshold.

The interaction of Arabidopsis with CH267 and *B. glumae* resulted also in large alterations in root metabolome and in the composition of root exudates. Camalexin is one of the most prominent metabolites playing a role in interaction with both pathogens and PGP bacteria (Koprivova and Kopriva, 2022) and correspondingly, camalexin was induced by both *B. glumae* and CH267, albeit to different level. The higher concentration of camalexin in roots, shoots, and exudates of plants treated with *B. glumae* compared to CH267 corresponds with the levels of induction of genes for camalexin biosynthetic enzymes as well as with increased protein accumulation of CYP71A12, CYP71A13, and GGP1 (Supplemental Figures S4, S8). Also the synthesis of GSH is induced on the gene expression, protein abundance, and metabolite accumulation levels, which is necessary for the increased camalexin synthesis but agrees also well with the induction of genes and proteins involved in GSH binding and GSH transferase activity. Another group of sulfur containing metabolites, the glucosinolates, which are also part of plant immune response, was only partly affected by the bacteria. Aliphatic glucosinolates were not affected, which was surprising, because they were previously associated with a defense specifically against bacterial pathogens (Fan et al., 2011). The indolic ones, however, were reduced in the shoots of plants treated with *B. glumae*, which might reflect the rerouting of the sulfur and indolic intermediates towards the synthesis of camalexin. However, the reduction also agrees with the upregulation of proteins for “glucosinolate catabolic processes” (Supplemental Table S5). Altogether, the alterations in accumulation of these compounds, and also cysteine (Figure 5) reflect the importance of sulfur in plant defense (Rausch and Wachter, 2005; Wang et al., 2022), which is supported by genes involved in sulfur homeostasis being among the upregulated DEGs after *B. glumae* treatment (Supplemental Table S2).

Among other metabolites, amino acids in the roots were highly affected by the treatment with both strains, more significantly with *B. glumae*. Indeed, several functional terms linked to amino acid metabolism were enriched in both transcriptome and proteome analysis of the roots, however, they were mostly connected to catabolism. This is corroborated by increased exudation of cadaverine derived from catabolism of lysine, and agrees with the emerging recognition of the importance of amino acid metabolism in plant microbe interactions (Moormann et al., 2022). However, while the amino acids were mostly increased in the roots, they were less abundant in the exudates, pointing to a possibility that the reduced exudation caused the accumulation in root tissues (Figure 5). In addition, in the proteome of roots treated with *B. glumae* many ribosomal proteins and proteins connected to translation were less abundant (Supplemental Table S5), which might also result in accumulation of amino acids. Myo-inositol was the metabolite most highly decreased in exudates from roots treated with both bacteria (Figure 5). This compound was shown previously to improve the colonisation ability of a PGP strain *Bacillus megaterium* YC4 (Vilchez et al., 2020). As its exudation is repressed by both CH267 and *B. glumae*, it seems that myo-inositol is not a universal signal but rather specific for certain bacterial taxa. On the other hand, indole 3-carboxylic acid is another indole besides camalexin that is increased in the exudates after bacterial treatments, corresponding to the upregulation of tryptophan metabolism on both transcriptome and proteome levels (Figures 4, 5). However, the detected changes in metabolites were not able to explain the opposite behaviour of Arabidopsis roots upon interaction with the two strains. It seems, therefore, that the impact of different strains is so multifaceted that a single time point analysis is not sufficient to fully understand the molecular mechanisms.

Beside the metabolites, the two strains affected the mineral composition of the roots, again in a different manner (Figure 6). Most remarkable difference is the accumulation of iron and zinc in roots treated with CH267, which corresponds with the induction of *IRT1* gene in the WT (Figure 2D, 6). Cluster 1 of the transcriptomic analysis also included gene for the key iron transporter IRT1. This is in accord with the recent finding of increased Fe uptake in Arabidopsis roots colonised by *Pseudomonas putida* (Esparza-Reynoso et al., 2024), and the general role of iron in plant immunity (Herlihy et al., 2020). The variation in accumulation of other elements, such as boron and magnesium reduced by both strains or potassium increased with CH267 and reduced with *B. glumae* (Figure 6), points to a possibility of more profound connection between mineral nutrition and plant microbe interactions, which is so far rather understudied.

### Effect of CYP71A27 on the interaction with CH267 and B. glumae

The multilevel analysis was also employed to help to dissect the function of CYP71A27. This gene was found in a GWAS for variation in microbiome activity, which resulted in the discovery of the new role of camalexin in interaction with PGP bacteria (Koprivova et al., 2019). Loss of CYP71A27 resulted in reduction in camalexin accumulation after treatments with CH267 (Supplemental Figure S4) but also to a loss of the growth promotion, which could be chemically complemented by camalexin (Koprivova et al., 2019). Interestingly, however, despite its sequence similarity, the CYP71A27 cannot complement the loss of the two enzymes for camalexin synthesis CYP71A12 and CYP71A13 (Figure 1C) and seems to be inactive. Together with its localization in phloem companion cells of the root this suggests that CYP71A27 must have a signalling function, possibly through protein-protein interactions. While the transcriptome and proteome analyses showed clearly that the *cyp71A27* mutant was more severely affected in interaction with the PGP strain than with the pathogen strain, based on the numbers of DEGs and DAPs, this was not true for the metabolites. On the metabolite level the impact of the mutation was seen on the sulfur containing camalexin and glutathione, and on few primary metabolites, mainly carbohydrates, but to a higher level in plants treated with *B. glumae*. The alterations in sugar accumulation are interesting while they might be connected to the specific phloem localization of CYP71A27 (Figure 1A, B) and its possible contribution to the shoot root communication in the control of camalexin synthesis (Koprivova et al., 2023). The relatively minor effects on metabolites compared to transcripts and proteins, corroborate the hypothesis that the CYP71A27 has a signalling and not metabolic function.

The important role of CYP71A27 in root shoot communication is apparent also from the proteomics data, since although the gene is almost not expressed in the shoots, its loss significantly affected the shoot proteome. Among the core 15 shoot proteins that are affected in *cyp71A27* and interacting with both bacteria (Figure 4D) are two proteins connected to xylem development, the PRX52 an apoplastic peroxidase involved in lignin polymerization (Floerl et al., 2012), as well as a xyloglucan endo-transglycosylase/hydrolase XTH4 that contributes to xylem fibre formation (Kushwah et al., 2020). In addition, the PMEI-PME25 is a pectin methyltransferase involved in cell wall modifications, as the PRX52. The regulation of proteins related to xylem in *cyp71A27* points to a possibility of a specific alterations in the vasculature. In addition, besides jasmonate and ABA, cytokinin signalling might be affected in *cyp71A27*, as the UDP-Glycosyltransferase UGT85A1 is involved in the homeostasis of trans-zeatin (Jin et al., 2013). Additional proteins significantly induced specifically in *cyp71A27* were connected to mitochondria in *B. glumae* treated plants, as e.g., the AT2G42310, the ESSS subunit of NADH:ubiquinone oxidoreductase (complex I) protein.

Indeed, mitochondria have important roles for plant immune response, including production of reactive oxygen species, release of cytochrome c, and initiation of programmed cell death (Wang et al., 2022). We observed increased accumulation of proteins for mitochondrial import and of components of both Complex I and Complex II (Supplemental Table S5) as well as stronger accumulation of several TCA cycle intermediates in the roots in *cyp71A27* (Figure 5). Possibly, the loss of CYP71A27 causes a disbalance in the immune response to the pathogenic *B. glumae*, where the response module of camalexin and other indoles is not affected, but the one connected with mitochondria is affected more. The mechanism how the loss of root localized CYP71A27 affects the leaf processes is, however, unclear.

In contrast to metabolites, which in *cyp71A27* are altered stronger by *B. glumae* than by CH267, the mineral composition of the mutant is affected more frequently by CH267 than by the pathogen (Figure 6). The lower accumulation of Fe compared to WT after CH267 treatment reflects the regulation of *IRT1* gene, i.e. high induction in WT than in *cyp71A27* (Figure 2D). The increased Fe uptake might be important for proper interaction with CH267, since the *irt1* mutant resembled *cyp71A27*, which is specifically affected by this strain.

Together with the regulation of genes for sulfur homeostasis this reveals an important role of plant microbe interaction for plant mineral nutrition and a function of CYP71A27 in mineral homeostasis during biotic interactions (Figure 6).

### New genes in plant microbe interaction network

To identify the genes most significantly contributing to the different response of *cyp71A27* to CH267 we firstly identified genes that were significantly transcriptionally affected under interaction of *cyp71A27* mutation and CH267 treatment (Figure 2D) and secondly, we performed a multilevel network analysis of the transcript and metabolite data to find genes directly correlating with the most significant metabolite changes in *cyp71A27* (Figure 7). The genes included in clusters 1 and 8 of the cyp27*CH267 dataset show the biggest difference between the mutant and WT specifically in CH267 treated roots (Figure 2D). The expression in *cyp71A27* and the similarity in response to CH267 of the mutants in cluster 1 genes *chitinase*, *cdef1*, *pr1-like*, and *irt1* with *cyp71A27* suggests that these genes contribute to the response of Arabidopsis to CH267. The phenotypes of the *cyp71A27* mutant thus seems to be caused indirectly, by the failed transcriptional regulation of these and other genes of the CYP71A27 transcriptional network. Interestingly, the response of *irt1* to CH267 is particularly weak, *CYP71B15* and *NAC042* are not induced at all and the *CYP71A12* is induced to a lower extent than in *cyp71A27*. This suggests a considerable importance of IRT1 for the interaction with CH267, corroborated by the changes in Fe content in the roots. Indeed, *IRT1* has been shown to be induced by several other plant growth promoting bacteria (Zamioudis et al., 2015; Orellana et al., 2022). Increased iron uptake might thus be one of the mechanisms for plant growth promotion, as iron is often limiting plant growth (Zamioudis et al., 2015). The lack of *IRT1* upregulation in *cyp71A27* after incubation with CH267 might thus be one of the mechanisms of the loss of growth promotion in this mutant.

However, also genes induced by CH267 only in the *cyp71A27* and not in the WT, i.e. genes from cluster 8, mimicked the *cyp71A27* phenotypes (Figure 3). The genes of the cluster were shown to be involved in defense against bacterial pathogens, such as nitrilase *NIT2* (Choi du et al., 2016), 3-deoxy-D-arabino-heptulosonate 7-phosphate synthase *DHS1* (Keith et al., 1991), vacuolar invertase *AT1G62660* (Bonfig et al., 2006), *RAP2.9* (Tsutsui et al., 2009), and *ANAC055* (Zheng et al., 2012). The asparagine synthase *ASN1* is interesting because another nitrogen connected gene, *NRT1;8*, was part of cluster 7 and because of the alterations in amino acid accumulation after inoculation of the roots with the two bacterial strains (Figure 5). Interestingly, several of these genes have also a function in modulating root architecture, such as the *NIT2* (Kutz et al., 2002), *ANAC055* (Lopez-Ruiz et al., 2022), and the vacuolar invertase (Sergeeva et al., 2006), which together with alteration in nitrate uptake and assimilation may point to the role of the cluster 8 genes in balancing the defense and root growth.

The lower camalexin accumulation in the mutants of cluster 1 and cluster 8 genes correlated with lower upregulation of *CYP71A12*, the key enzyme in camalexin synthesis in the roots. This is not surprising for *anac055* mutant, because *ANAC055* encodes for a transcription factor in stress related networks, such as a network of response to *Botrytis cinerea*, in which it controls downstream factors, including WRKY33, a direct regulator of *CYP71A12* (Hickman et al., 2013; Nguyen et al., 2022). Also *RAP2.9* encodes a transcription factor, a member of the ethylene responsive ERF family, and since camalexin is controlled also by ethylene signalling, it is possibly also involved in control of camalexin synthesis (Zhou et al., 2022). The mechanism(s) linking the loss of other tested genes with low induction of *CYP71A12* and lower camalexin accumulation, however, remain elusive.

An independent approach to identify genes highly connected to the function of CYP71A27 in response to CH267 is a multilevel network analysis. These networks make use of the availability of omics datasets and can be composed from different data types (Garcia-Gomez et al., 2020). The advantage of combining gene co-expression networks with e.g. metabolite accumulation has been shown multiple times, e.g. by using such networks of sulfur starvation response to identify genes encoding sulfotransferases in glucosinolate synthesis (Hirai et al., 2005). In our study the networks were generated for each condition separately, i.e. the main *cyp71A27* effect and the interactions with CH267 and *B. glumae*, to better dissect the specific functions of CYP71A27 in different conditions. Since the smallest network, i.e. the least significant correlations, were observed after treatment with *B. glumae* (Figure 7, Supplemental Figure S13), the loss of CYP71A27 seems to have a relatively small effect on the outcome of the interaction. This is reflected also in the number of genes differently regulated in *cyp71A27* and the WT, which is much smaller in *B. glumae* treated roots than in the WT (Figure 2). As already suggested from the general transcriptome alterations, the most correlations were found in the DEGs from interaction with CH267. One of the major hub genes, the *LTPG30*, has a similar expression pattern in the root vasculature as *CYP71A27* (van der Graaff et al., 2002). Overexpression of *LTPG30*, also named as *VASCULAR TISSUE SIZE* (*VAS*), resulted in increased number of vascular cells (van der Graaff et al., 2002), which might be connected with several genes for cell wall modification being found in the Cluster 1 of DEGs from interaction between *cyp71A27* and CH267 effects. The other major hub gene, *YSL9*, encodes an uncharacterised plastidial protein, and is induced during interaction with bacteria and fungi. Interestingly, some genes of the network directly connected to camalexin or GSH are co-localised with CYP71A27 in the root vasculature, but some, such as the defense related *YSL9*, *CHITINASE*, *BSP*, or *PER23* are not, thus they seem to be connected rather through the metabolic network downstream the action of CYP71A27 in response to CH267.

This is corroborated by the analyses of the four mutants in genes from the *cyp71A27*\*CH267 network, which as before, showed the same low accumulation of camalexin and low upregulation of the marker genes as the *cyp71A27* mutant (Figure 8). The *bsp* mutant is particularly interesting in this context, because it has low camalexin but the genes for its synthesis *CYP71A12* and *CYP71B15* are regulated like in WT (Figure 8). This points to a posttranscriptional regulation of camalexin synthesis, which might be connected to the metabolome arrangement of the biosynthetic enzymes described for Arabidopsis leaves (Mucha et al., 2019). Another interesting result is the interconnection of camalexin synthesis with biosynthesis of thalianol. This triterpene has been shown to modulate the composition of Arabidopsis root microbiome (Huang et al., 2019), so the interplay of these two secondary metabolites may allow a more tailored response to diverse microbiota in the root rhizosphere. Finding BHLH39 in the camalexin network and the phenotype of the corresponding mutant (Figure 8) aligns well with the changes in *IRT1* expression, phenotype of *irt1* mutant, and differences in accumulation of iron, because this transcription factor belongs to the regulatory network controlling iron uptake and homeostasis (Wang et al., 2007; Cui et al., 2018). BHLH39 is also linked to jasmonate and ethylene signalling (Cui et al., 2018; Yang et al., 2022), connecting the genes from network analysis with those from cluster 1 and cluster 8 derived from the transcriptomics. The analysis of the *bhlh39* mutant further corroborates the importance of iron for the interaction of Arabidopsis with CH267.

**Figure 8.**
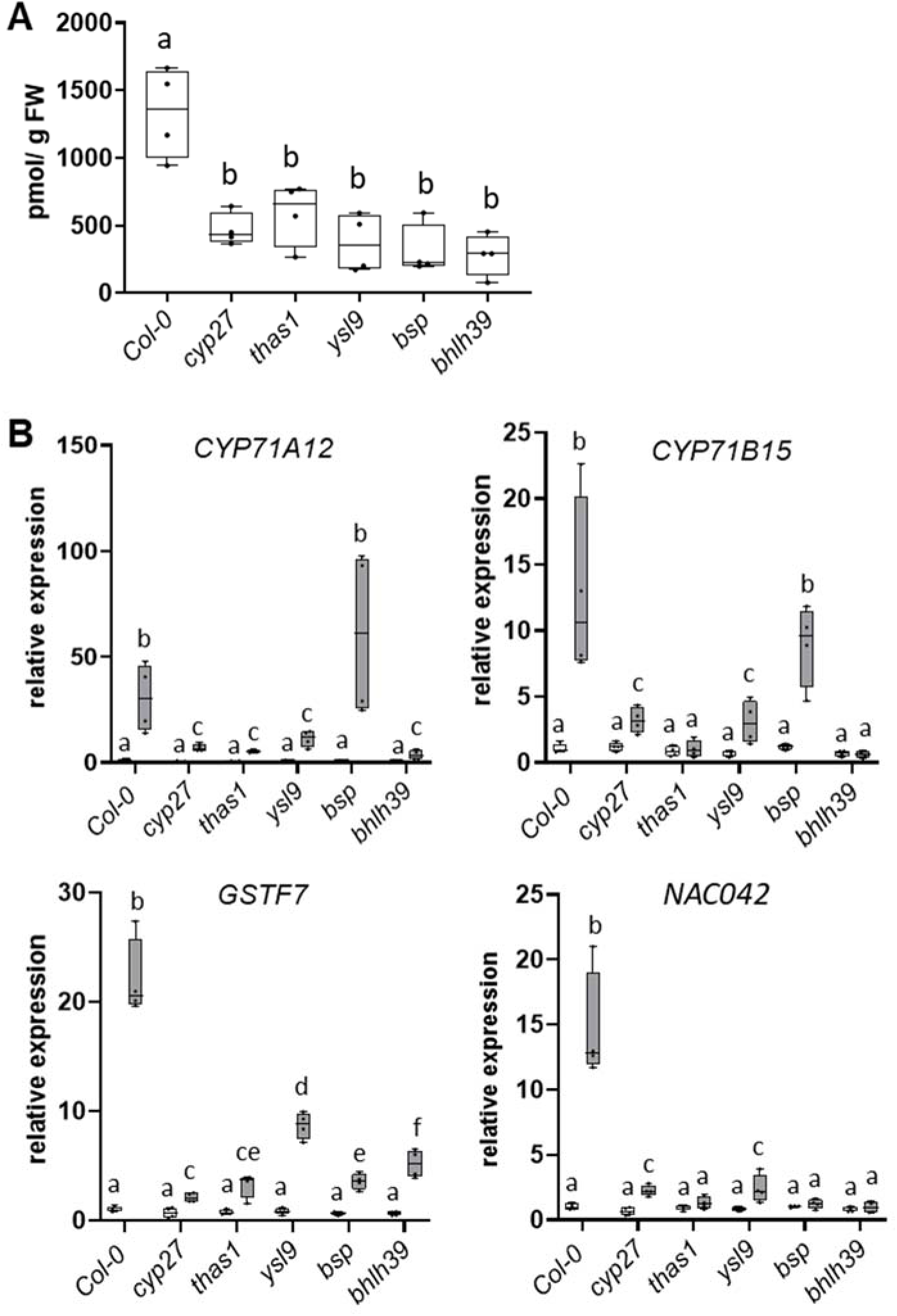
Analysis of mutants in candidate genes from network analysis. **A** Camalexin levels in shoots of WT, *cyp71A27*, and mutants in candidate genes from network analysis – *thas1, ysl9, bsp,* and *bhlh39* treated with CH267. **B** Relative transcript levels of marker genes responding to bacterial infection - *CYP71A12, CYP71B15, GSTF7, NAC042* – in roots of WT, *cyp71A27*, *thas1, ysl9, bsp,* and *bhlh39* treated with mock or CH267. The qRT-PCR results were obtained from 4 biological and 2 technical replicates, the transcript levels of WT Col-0 in mock treated roots was set to 1. Letters mark values significantly different at P<0.05 (ANOVA).

Altogether, here we show that CYP71A27 is a non-canonical P-450 without a metabolic function but active in signalling as its loss affects the transcriptome more than a metabolome. Correspondingly, immunity and developmental processes seem to be more affected in *cyp71A27* mutant than metabolism, maybe with the exception of the camalexin accumulation. The multifaceted signalling function is evidenced by the variety of mutants of genes affected by the loss of CYP71A27 having similar response to CH267 as the *cyp71A27* mutant. This is true for genes upregulated more strongly in the *cyp71A27* than in WT upon treatment with CH267 as well as those repressed specifically in the CH267 treated mutant. The genes analysed here were not connected to interaction with PGP strains before and represent thus a valuable resource to better understand the complex interplay between plants and commensal microorganisms. The CYP71A27 itself seems to be an important hub in this interplay.

## MATERIALS AND METHODS

### Plant material and growth conditions

*Arabidopsis thaliana L.* accession Col-0 was used as wild type throughout. The mutant *cyp71A27* (SALK_053817) was described previously (Koprivova *et al*., 2019). The mutants *chitinase* (Salk_043790)*, cdef1* (SALK_014093)*, pr1-like* (SALK_014249C)*, irt1* (SALK_097869C)*, anac055* (SALK_011069C), *rap2.9* (SALK_052297), *nit2* (SAIL_681_H09), *asn1* (GK-829B05) *thas1* (SAIL_779_D03)*, ysl9* (SAIL_91_D09)*, bsp* (SALK_128725), and *bhlh39* (SALK_025676C) were obtained from Nottingham Arabidopsis Stock Centre (NASC) and genotyped by PCR to recover homozygous lines (Alonso et al., 2003). The *cyp71A12 cyp71A13* (Muller et al., 2015) mutant was obtained from Dr. H. Frerigmann, MPIPZ Cologne.

The plants were grown in a hydroculture system as described previously (Koprivova et al., 2023). Seeds were surface sterilized with chlorine gas, resuspended in 0.1 % agarose, distributed onto square 1 cm x 1 cm sterile nylon membranes (ca. 30 seeds per sample) and placed in 12 well plates on top of 1 ml of ½ Murashige Skoog (MS) medium with 0.5 % sucrose. The plates were kept for 2 days in 4°C and dark for stratification and then transferred to 22°C and in dark for further 3 days to promote etiolation. Afterwards the plates were incubated in controlled environment at long day conditions (16 h light/ 8 h dark), 120 µE m^-2^ S^-1^, and at 22°C for 7 days. The medium was replaced with ½ MS without sucrose and the plants were incubated for 24 hours before inoculation with the bacteria or mock and incubated further in the same conditions for 3 days.

### Bacterial strains and cocultivation conditions

For bacterial treatments *Pseudomonas* sp. CH267 (Haney *et al*., 2015), obtained from J. R. Dinneny, Stanford University and *Burkholderia glumae* PG1 (Gao *et al*., 2015), obtained from K.-E. Jäger, Heinrich Heine University Düsseldorf, Germany, were used. The bacteria were kept as glycerol stocks and plated freshly before experiment on LB plates supplemented with appropriate antibiotics grown at 30°C. For inoculation, overnight bacterial cultures were washed two times with sterile 10 mM MgCl_2_ and diluted to OD_600_ = 0.0001 (CH267) to or OD_600_ = 0.0005 (*B. glumae*). Each well of the 12-well plate was inoculated with 8 µl of the bacterial suspension or 10 mM MgCl_2_ as mock. Samples for were harvested after 3 days of inoculation.

### Functional analysis of CYP71A27

For complementation of the *cyp71A12 cyp71A13* mutant, genomic fragments containing the coding regions of CYP71A12 and CYP71A27 were amplified by PCR with primers AK12N (caccATGGAGATGATATTGATGGTC)/AK12C (AAAAAGAGAAACATTTTAAATAACG) and AK27N (caccATGGAGATGATATTAATCTCTTTG)/AK27C (GTTTTCTCATCAATGTGTGTGC), respectively, and cloned into pENTR/D-TOPO vector (Invitrogen). After verification of the sequence by Sanger sequencing the fragments were transferred to destination vector pAMPAT-35S-GW (Pesch et al., 2015) by LR. The resulting plasmid was transformed to *Agrobacterium tumefaciens* GV3101 (pMP90) and *cyp71A12 cyp71A13* plants were transformed by the floral dip method and the transformants were selected by BASTA.

Four independent transgenic lines per construct were grown in the 12 well plates, treated with *B. glumae* for 3 days as described before and camalexin was measured in the leaves. To verify expression of the transgene, RNA was isolated from the roots and the transcript levels of *CYP71A12* or *CYP71A27* were determined by qRT-PCR.

### Microscopy

The CYP71A27 was translationally fused with GFP after amplification of the genomic fragment by PCR with primers AK27N /AK27F (GTTTTCTGATCAATGTGTGTGC), cloning into pENTR/D-TOPO, and transferring to destination vector pAMPAT-35S-GW (Pesch et al., 2015) by LR. Afterwards the 35S promoter was exchanged by a promoter of CYP71A27, which was amplified from genomic DNA by primers CYP27PROASC (atatGGCGCGCCtttgacctatactttactacc) and CYP27PROXho (atatCTCGAGattgtgcttatttaggaatgg) and cloned into pCR2.1 TOPO plasmid. After sequencing the promoters were excised by XhoI and AscI restriction enzymes and the CYP71A27 promoter was inserted in the cut pAMPAT-CYP27::GFP vector. The resulting plasmid was transformed to *Agrobacterium tumefaciens* GV3101 (pMP90) and *cyp71A27* plants were transformed by the floral dip method and the transformants were selected by BASTA.

Imaging was performed on a Zeiss LSM 980 Airyscan 2 system equipped with a 63x (NA1.4) oil immersion lens or a 20x (NA 0.8) air lens. All plants were fixed and cleared according to a previously described protocol (Ursache et al., 2018). Cell walls were stained using Calcoflour white CW). CW signals was collected by excitation with a 405 nm laser and emission collected between 420-435 nm). For GFP-signal, excitation was 488 nm and emission collected in a 510-545 nm window.

### Camalexin measurements

Camalexin was extracted from approx. 10-20 mg of plant roots or shoots and determined by HPLC as described in (Koprivova et al., 2019). For determination of camalexin in the exudates the procedure described in (Koprivova et al., 2023) was used. The nutrient solution was centrifuged and purified using 1 ml solid phase extraction tubes (Discovery-DSC18).

After elution with 90% (V/V) acetonitrile and 0.1% (V/V) formic acid, the camalexin was determined by HPLC. For the quantification external standards were used ranging from 1 pg to 1 ng per µl.

### RNA isolation and quantitative RT-PCR

Total RNA was isolated from roots by standard phenol / chlorophorm extraction and LiCl precipitation. First strand cDNA synthesis was performed using QuantiTect Reverse transcription Kit (Qiagen) from 800 ng of total RNA. Quantitative real time RT-PCR (qPCR) was performed using gene-specific primers (Supplemental Table S10) and the fluorescent dye SYBR Green (Promega) as described in (Koprivova et al., 2023). All quantifications were normalized to the *TIP41* (AT4G34270) and *UBC21* (AT5G25760) genes. The RT-PCR reactions were performed in duplicate for each of the 4 independent samples.

### RNA sequencing

For RNA sequencing the RNA was isolated with PureLink™ Plant RNA Reagent (Invitrogen, Thermo Scientific™, Waltham, Massachusetts, USA), using the small-scale isolation procedure as described in (Zenzen et al., 2024). The RNA samples were further treated with TURBO DNA-free™ Kit (Invitrogen, Thermo Scientific) to eliminate DNA, according to the manufactureŕs protocol. Purified RNA was sequenced by Novogene (Novogene Co., Ltd., Cambridge, UK) using a non-strand-specific RNA library construction by poly-T mRNA enrichment, random fragmentation, and cDNA synthesis via random hexamer priming. The RNA was sequenced using Illumina NovaSeq™ 6000 Sequencing System (Illumina, Inc., San Diego, California, USA), to produce paired-end 150 bp reads and coverage of 30-40 Mio reads per sample. The adapters from the raw reads were removed by processing with Trimmomatic (Bolger et al., 2014) and the reads were then mapped to the *A. thaliana* reference genome (TAIR10) using HiSAT2 with default parameters (Kim et al., 2019). The resulting reads were then counted with HTseq (Anders et al., 2015) with default parameters and normalised to TPM values. The differential expression analysis was performed with DESeq2 (Love et al., 2014) to obtain the main effect of *cyp27* genotype and the interaction DEGs for G*E (cyp27*CH or cyp27*BG) and limma (Ritchie et al., 2015) to obtain the effect of CH and BG treatments in the two genotypes. We used log2 fold change >1 or log2 fold change <-1, and adjusted P-value <0.01 cut off to obtain DEGs for further analyses. The Venny platform (https://bioinfogp.cnb.csic.es/tools/Venny/index.HTML) was used to construct the Venn diagrams. Z-score was computed on a trait-by-trait (e.g. gene-by-gene) basis by subtracting the mean and then dividing by the standard deviation. The obtained Z score was further used to create the heatmaps. Clustering was performed using the MeV software (http://mev.tm4.org/). Gene ontology (GO) and KEGG pathway enrichment analyses were performed in Biomaps app in VirtualPlant (Katari et al., 2010). The graphs were constructed using the SRplot platform (Tang et al., 2023). The RNAseq data have been submitted to the NCBI SRA repository under accession number PRJNA1196130.

### Proteomics

The proteomics analysis was performed exactly as in (Berkova et al., 2024). Briefly, approximately 10 mg of homogenized freeze-dried plant material was extracted with methyl tert-butyl ether/methanol/water mixture. Proteomics samples corresponding to 5 µg of proteins were analyzed by nanoflow reverse-phase liquid chromatography-mass spectrometry on a 15 cm C18 Zorbax column (Agilent), a Dionex Ultimate 3000 RSLC nano-UPLC system, and the Orbitrap Fusion Lumos Tribrid Mass Spectrometer (Thermo Fisher Scientific). The measured spectra were recalibrated and searched against the *A. thaliana* ARAPORT 11 database and common contaminants databases using Proteome Discoverer 2.5 (Thermo Fisher Scientific) with algorithms SEQUEST and MS Amanda (Dorfer et al., 2014). The quantitative differences were determined by Minora, employing precursor ion quantification followed by normalization (total area) and calculation of relative peptide/protein abundances. Only proteins with at least two unique peptides were considered. Protein function was predicted by searching sequences against the *A. thaliana* proteome in STRING 11.0 and UniProt (https://www.uniprot.org/). The analysis was done in at least four biological replicates. The mass spectrometry proteomics data were deposited at the ProteomeXchange Consortium via the PRIDE partner repository (Perez-Riverol et al., 2022) with the data set identifier PXD058537. Statistical evaluation was performed as described for the RNA sequencing dataset, using an absolute log2 fold change >1 and an adjusted p-value <0.01 as cutoffs to identify the most relevant DAPs for further analyses. Only proteins with more than 4 values out of the five replicates in at least one condition were considered for the analysis.

### Metabolite analysis

Thiols and glucosinolates were determined by standard HPLC procedures as described in (Dietzen et al., 2020). The untargeted analysis of root metabolites was performed as described in (Zenzen et al., 2024) from ca. 30 mg frozen plant material. The metabolites were derivatized using methoxyamine hydrochloride and N-Methyl-N-(trimethylsilyl)trifluoroacetamide (MSTFA) and analyzed by gas chromatography coupled to mass spectrometry as described (Shim et al., 2019). The identification was performed using the MassHunter Workstation Qualitative Analysis Software (version B.06.00, Agilent Technologies, Santa Clara, California, USA) by comparing the spectra to the NIST14 Mass Spectral Library (https://www.nist.gov/srd/nist-standard-reference-database-1a-v14). Quantitative measurement was achieved by peak integration with MassHunter Workstation Quantitative Analysis Software (version B.08.00), using the internal standard ribitol (Zenzen et al., 2024).

The exudates were freeze dried in SpeedVac. The exudates samples were subsequently derivatized as described in (Berkova et al., 2024) and analyzed using a Q Exactive GC Orbitrap GC-MS/MS mass spectrometer (Thermo Fisher Scientific) coupled to a Trace 1300 Gas chromatograph (Thermo Fisher Scientific) using a TG-5SILMS column (30 m; 0.25 mm; 0.25 μm; Thermo Fisher Scientific) with a temperature gradient (5 min at 70 °C followed by a 9 °C gradient in 1 min to 320 °C and final incubation for 5 min at 320 °C). Data were analysed by Compound Discoverer 3.3 (Thermo) and searched against NIST2014, GC-Orbitrap Metabolomics library, and in-house library. Only metabolites that fulfilled identification criteria (score ≥ 85 and ΔRI < 1%) were included in the final list of identified compounds. The quantitative differences were validated by manual peak assignment in Skyline 19.1 (Pino et al., 2020), using the extracted ion chromatogram (2 ppm tolerance). The analysis was done in four biological replicates.

### Elemental analysis

The mineral composition of leaf tissue was determined by inductively coupled plasma mass spectrometry (ICP-MS) from 15-20 mg lyophilized shoot tissue by the CEPLAS Plant Metabolism and Metabolomics Laboratory, University of Cologne, using an Agilent 7700 ICP-MS (Agilent Technologies, Santa Clara, CA, USA) (Almario et al., 2017).

### Network analysis

Multivariate networks between the DEGs and metabolites were created in Cytoscape (Shannon et al., 2003) using the significant (*P*-value <0.05) pair-wise correlations of DEGs and metabolite levels. The Pearson correlation analysis was performed using the ‘Hmisc’ and ‘corrplot’ packages in R (https://www.R-project.org).

## Data availability

The RNAseq data are available through the NCBI SRA repository under accession number PRJNA1196130. The mass spectrometry proteomics data were deposited at the ProteomeXchange Consortium with the data set identifier PXD058537.

## Additional Information

**Supplemental Figure S1.** Transcriptome response of Arabidopsis roots to bacteria.

**Supplemental Figure S2.** Enrichment analysis of the transcriptome response of Arabidopsis roots to bacteria.

**Supplemental Figure S3.** Effect of loss of CYP71A27 on transcriptome response of Arabidopsis roots to bacteria.

**Supplemental Figure S4.** Camalexin synthesis after treatment with *B. glumae* PG1 (BG) and *Pseudomonas* sp. CH267 (CH).

**Supplemental Figure S5.** Expression of the candidate genes in roots of *cyp71A27* and WT after treatments with mock, CH267, and *B. glumae*.

**Supplemental Figure S6.** Effect of loss of *CYP71A27* on the proteome response of Arabidopsis to bacteria.

**Supplemental Figure S7.** Enrichment analysis of the proteome response of Arabidopsis roots and shoots to bacteria.

**Supplemental Figure S8.** Pathway of camalexin synthesis and heat map with corresponding protein abundance.

**Supplemental Figure S9.** Comparison of root and shoot DAPs in WT Col-0 and *cyp71A27*.

**Supplemental Figure S10.** Comparison of DEGs and DAPs in roots of WT Col-0 and *cyp71A27* after treatments with CH267 or *B. glumae*.

**Supplemental Figure S11.** Metabolite analysis of response of Arabidopsis to *Pseudomonas* sp. CH267 and *Burkholderia glumae* PG1.

**Supplemental Figure S12.** Camalexin in exudates of Arabidopsis treated with *Pseudomonas* sp. CH267 and *Burkholderia glumae* PG1.

**Supplemental Figure S13.** Network analysis of DEGs and metabolites.

**Supplemental Table S1.** Differentially expressed genes in *cyp71A27* and WT Arabidopsis roots after treatment with *Pseudomonas* sp. CH267 and *Burkholderia glumae* PG1.

**Supplemental Table S2.** Enrichment analyses of GO terms in DEGs from Arabidopsis roots of *cyp71A27* mutant and WT Col-0 after treatment with *Pseudomonas* sp. CH267 and *Burkholderia glumae* PG1.

**Supplemental Table S3.** Enrichment analyses of KEGG pathways in DEGs from Arabidopsis roots of *cyp71A27* mutant and WT Col-0 after treatment with *Pseudomonas* sp. CH267 and *Burkholderia glumae* PG1.

**Supplemental Table S4.** Complete proteomics data for roots and shoots.

**Supplemental Table S5.** Enrichment analyses of GO terms in DAPs from Arabidopsis roots and shoots of *cyp71A27* mutant and WT Col-0 after treatment with *Pseudomonas* sp. CH267 and *Burkholderia glumae* PG1.

**Supplemental Table S6.** Enrichment analyses of KEGG pathways in DAPs from roots and shoots of *cyp71A27* mutant and WT Col-0 after treatment with *Pseudomonas* sp. CH267 and *Burkholderia glumae* PG1.

**Supplemental Table S7.** Comparison of DEGs and DAPs in Arabidopsis Col-0 and *cyp71A27* roots after treatment with *Pseudomonas* sp. CH267 and *Burkholderia glumae* PG1.

**Supplemental Table S8.** Metabolite analysis of roots and exudates of Arabidopsis Col-0 and *cyp71A27* roots after treatment with *Pseudomonas* sp. CH267 and *Burkholderia glumae* PG1.

**Supplemental Table S9.** Genes directly connected to the 8 selected metabolites in multilevel network analysis.

**Supplemental Table S10.** Primers used in the study.

## Acknowledgements

We thank I. Klinkhammer, B. Welter and S. Ambrosius for technical support. We also thank the Biocenter MS Platform Cologne for the measurements of mineral composition and the CEPLAS metabolism & metabolomics laboratory for the analysis of primary metabolites. We thank Dr. M. Jacoby (University of Cologne) for the pAMPAT-35S-GW plasmid and Dr. H. Frerigmann (MPI-PZ Cologne) for the *cyp71A12 cyp71A13* seeds. G.M.T. thanks for support from the International Max Planck Research School on “Understanding Complex Plant Traits Using Computational and Evolutionary Approaches” and from the German Academic Exchange Service (DAAD) - project 57655738.

## Author contributions

S.K. and A.K. conceived and designed the research, A.K., M.B., V.B., G.M.T., T.G.A., and P.W. performed the experiments. D.R. performed most of the bioinformatics analysis, M.Č. was involved in proteomics analysis. S.K. drafted the manuscript, A.K., D.R., M.B. and S.K. prepared figures. All authors were involved in the revising of manuscript.

