## supplementary figures for "Multilevel analysis of response to plant growth promoting and pathogenic bacteria in Arabidopsis roots and the role of CYP71A27 in this response"

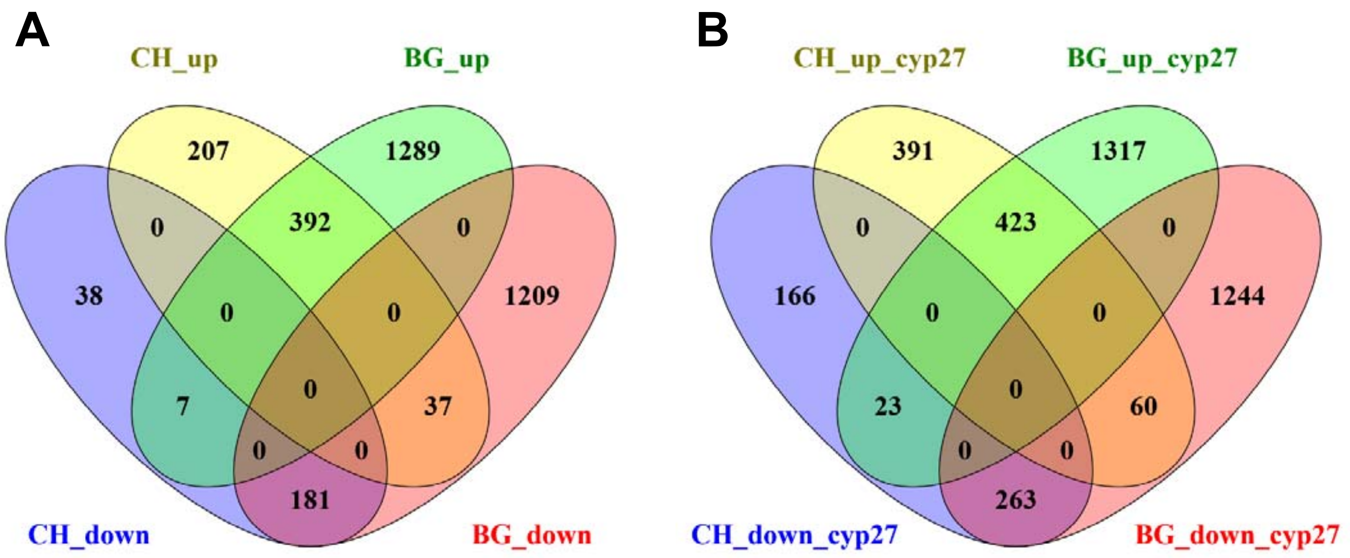


**Supplemental Figure S1.** Transcriptome response of Arabidopsis roots to bacteria. **A** Venn diagram showng the number of DEGs in WT roots treated with *Pseudomonas* sp. CH267 (CH) and *Burkholderia glumae* PG1 (BG) compared to mock treatment. **B**. Venn diagram showng the number of DEGs in *cyp71A27* roots treated with *Pseudomonas* sp. CH267 (CH) and *Burkholderia glumae* PG1 (BG) compared to mock treatment.

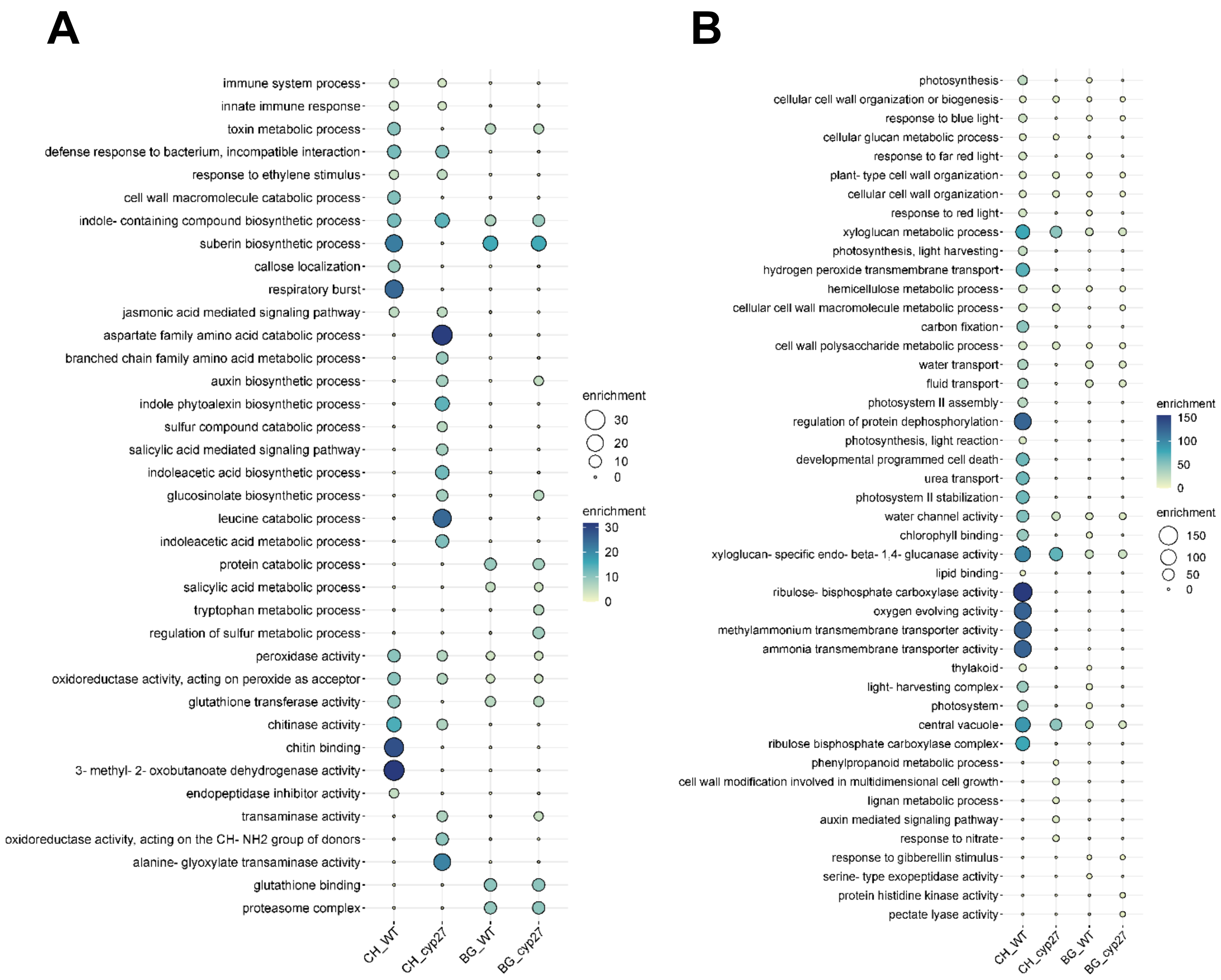


**Supplemental Figure S2.** Enrichment analysis of the transcriptome response of Arabidopsis roots to bacteria. The bubble plots show GO functional categories overrepresented among upregulated (**A**) and downregulated (**B**) DEGs after treatment of *cyp71A27* (cyp27) and WT with Pseudomonas CH267 (CH) and *B. glumae* (BG). The size and colour of the circles correspond to the enrichment.

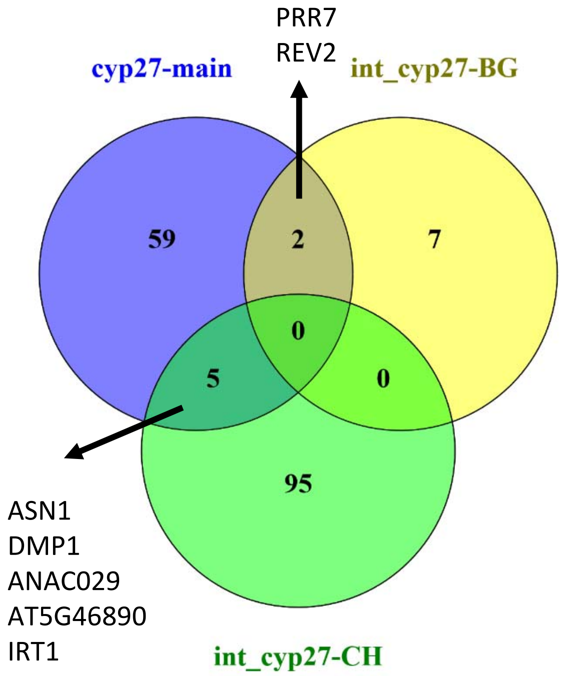


**Supplemental Figure S3.** Effect of loss of CYP71A27 on transcriptome response of Arabidopsis roots to bacteria. Venn diagram showing the number of DEGs in *cyp71A27* main effect, and interaction of cyp27 with CH and BG.

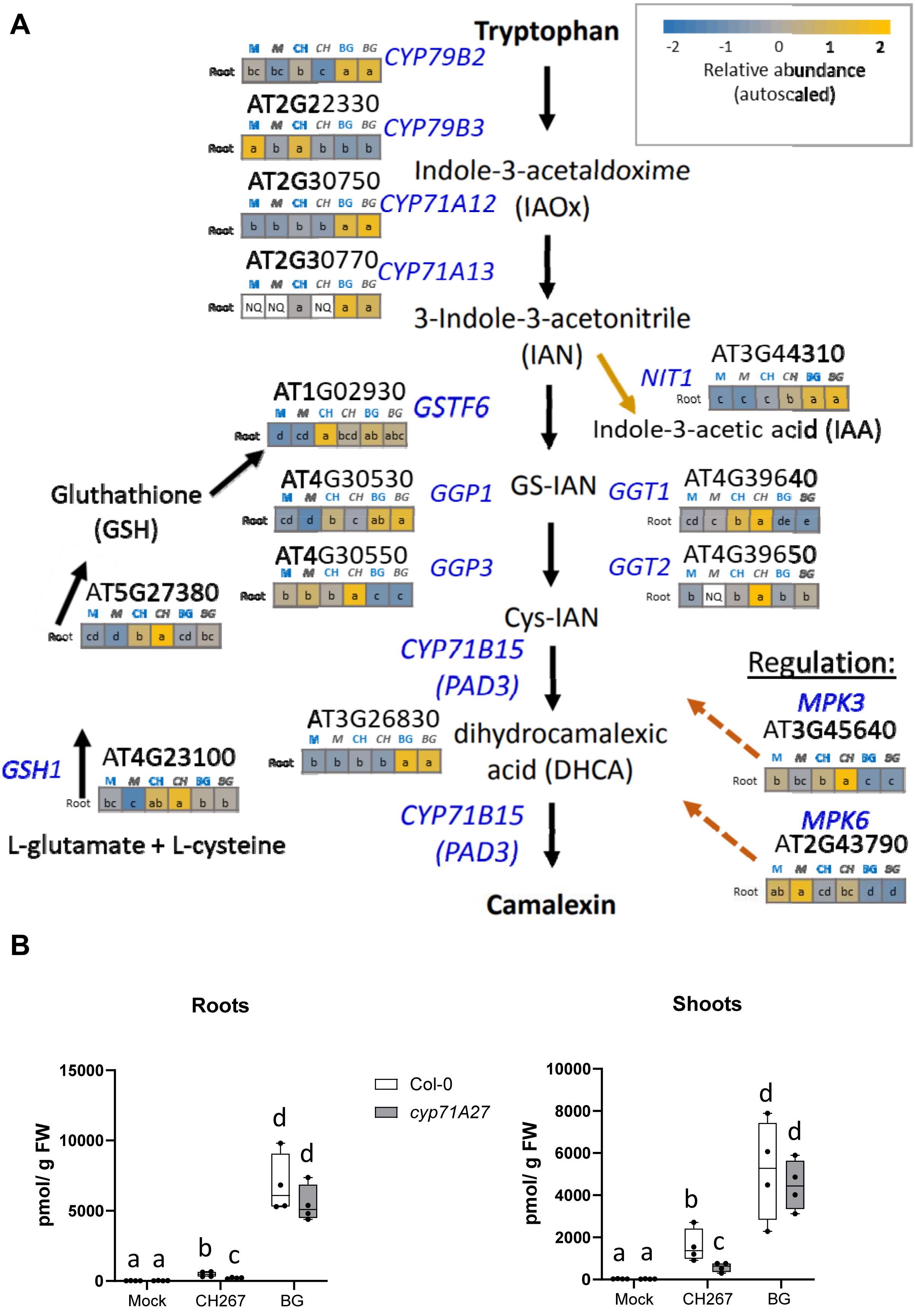


**Supplemental Figure S4**. Camalexin synthesis after treatment with *B. glumae* PG1 (BG) and *Pseudomonas* sp. CH267 (CH). (A) Pathway of camalexin synthesis and heat map with correspnding gene expression. (B) Camalexin concentration in roots and shoots determined by HPLC. Letters mark significantly different values at $p<0.05$ (Student T-test).

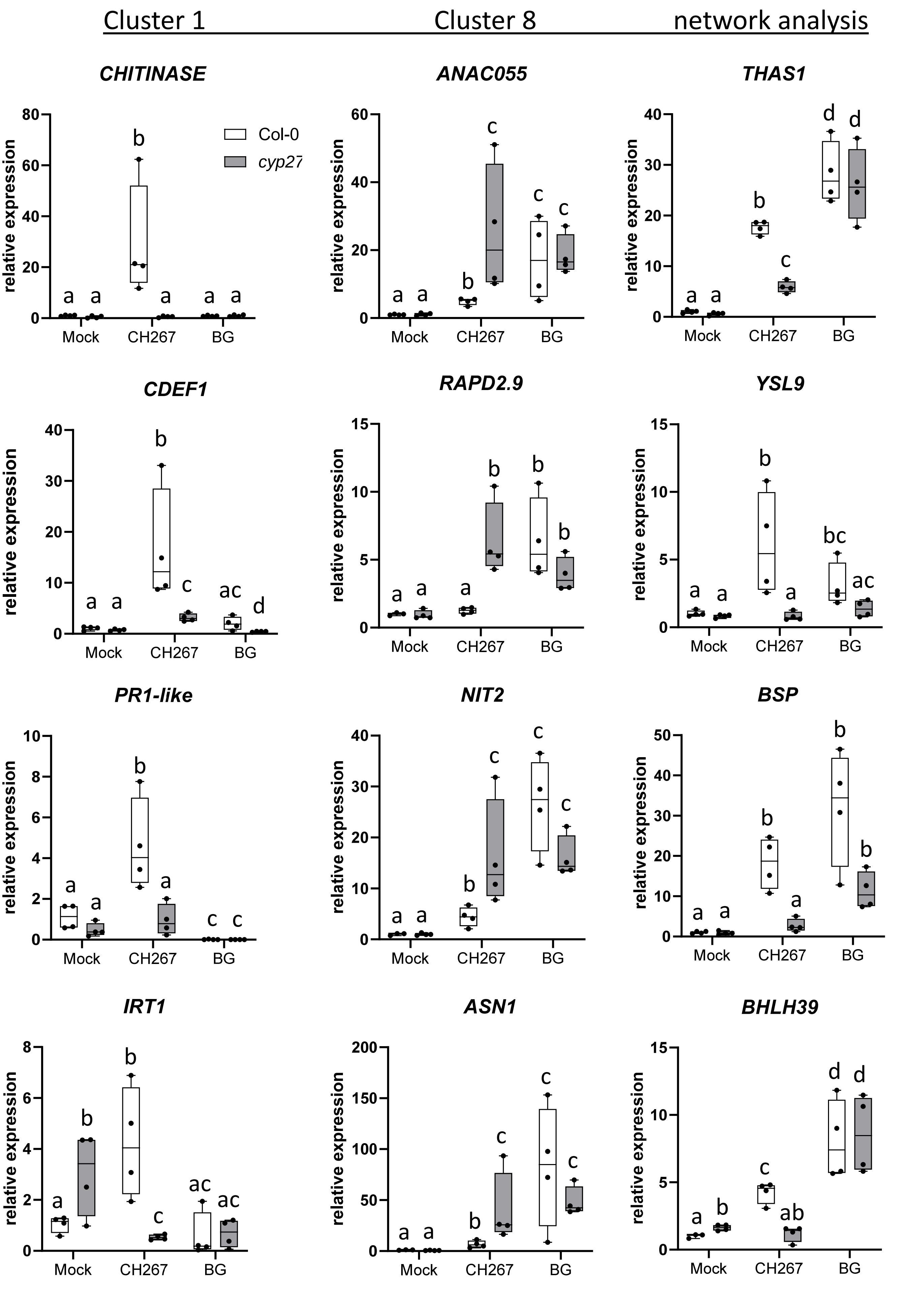


**Supplemental Figure S5.** Expression of the candidate genes in roots of *cyp71A27* (grey) and WT (white) after treatments with mock, CH267, and *B. glumae* (BG) determined by qRT-PCR. Different letters mark values significantly different at $p<0.05$ (T-test).

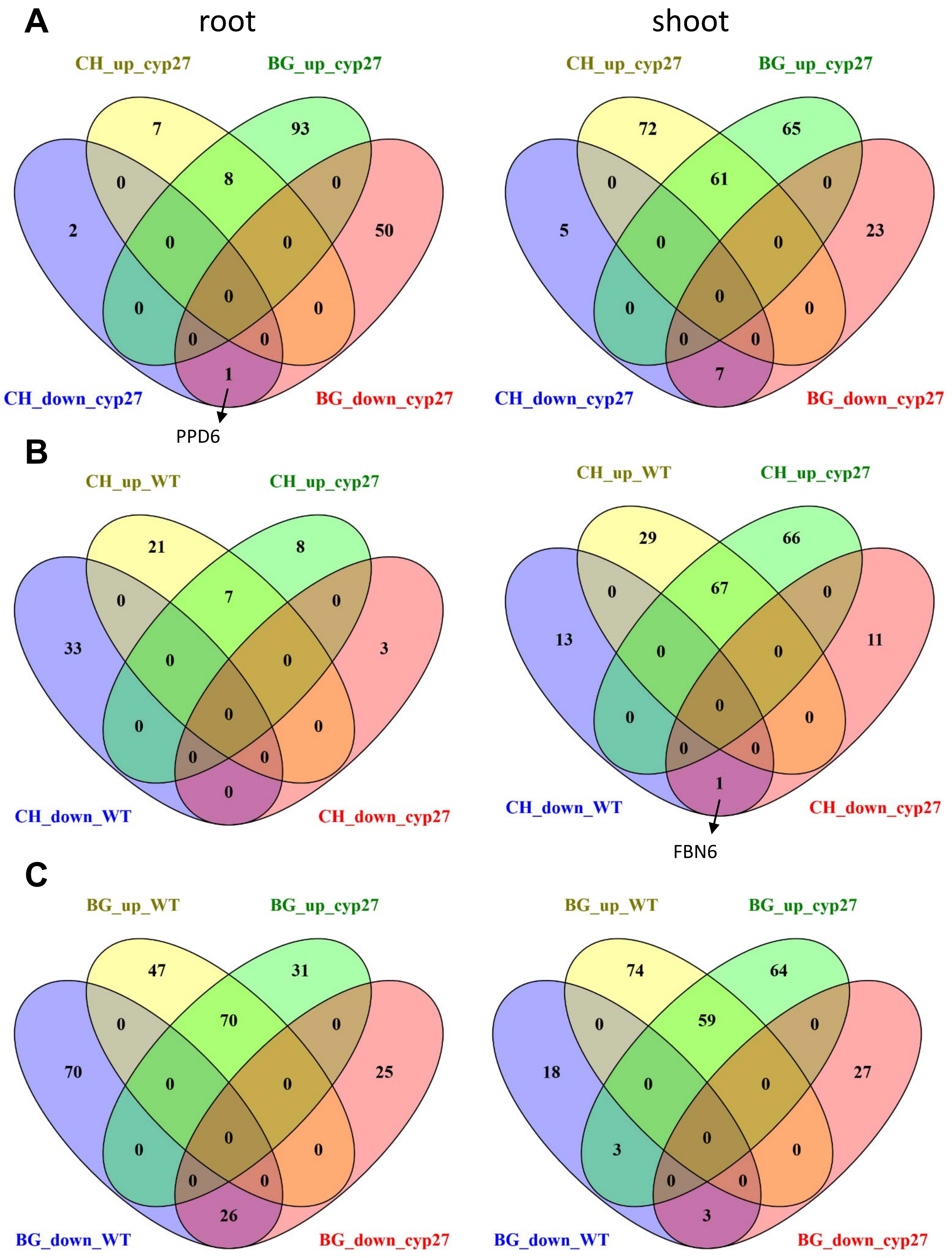


**Supplemental Figure S6.** Effect of loss of *CYP71A27* on the proteome response of Arabidopsis to bacteria. **A** Venn diagram showng the number of DAPs in *cyp71A27* roots and shoots treated with *Pseudomonas* sp. CH267 (CH) and *Burkholderia glumae* PG1 (BG) compared to mock treatment. **B**. Venn diagram comparing the number of DAPs regulated by CH267 vs. mock in *cyp71A27* mutant and WT **C** Venn diagram comparing the number of DAPs regulated by *B. glumae* vs. mock in *cyp71A27* mutant and WT

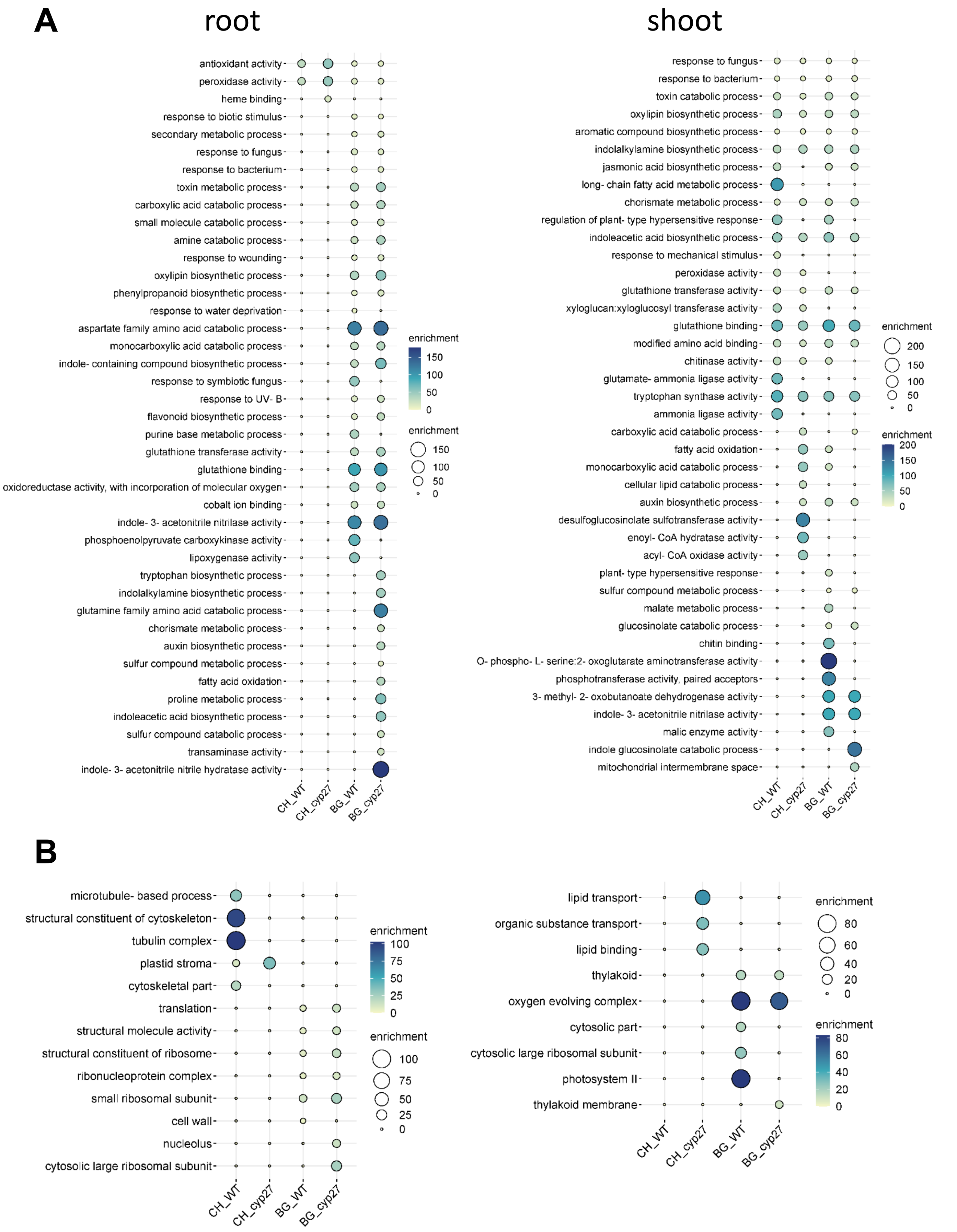


**Supplemental Figure S7.** Enrichment analysis of the proteome response of Arabidopsis roots and shoots to bacteria. The bubble plots show GO functional categories overrepresented among upregulated (**A**) and downregulated (**B**) DEGs after treatment of *cyp71A27* (cyp27) and WT with Pseudomonas CH267 (CH) and *B. glumae* (BG). The size and colour of the circles correspond to the enrichment.

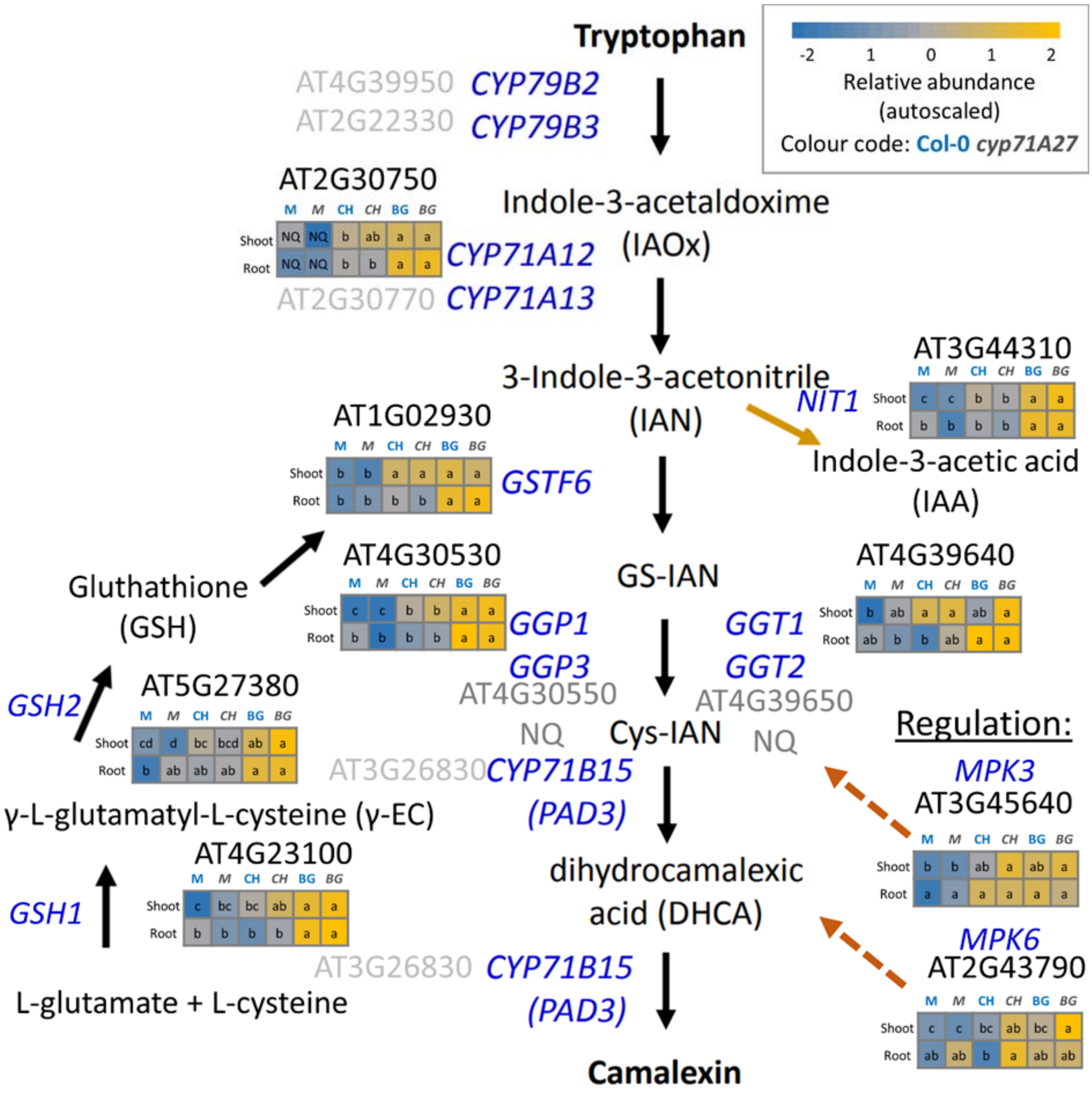


**Supplemental Figure S8.** Pathway of camalexin synthesis and heat map with corresponding protein abundance.

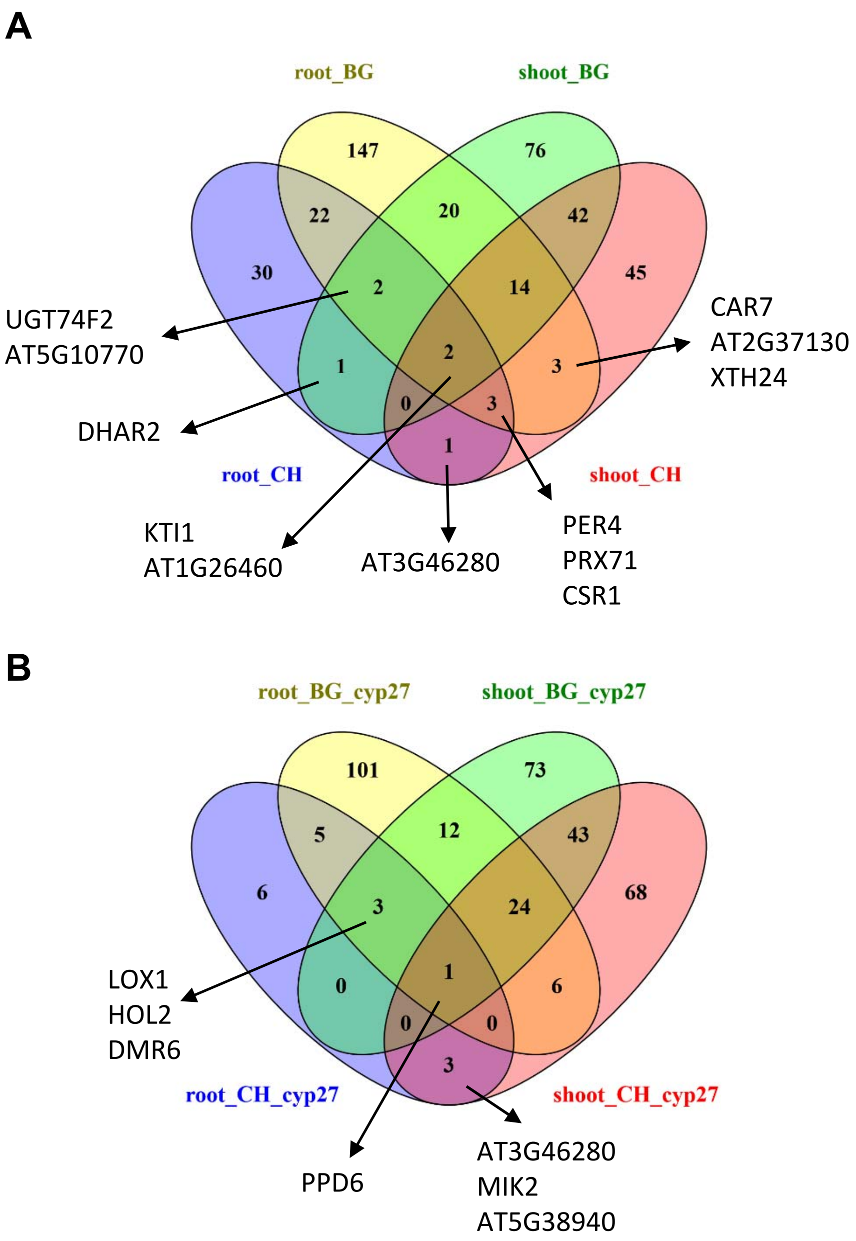


**Supplemental Figure S9.** Venn diagram showing comparison of root and shoot DAPs in WT Col-0 (A) and *cyp71A27* (**B**), with some interesting DAPs indicated.

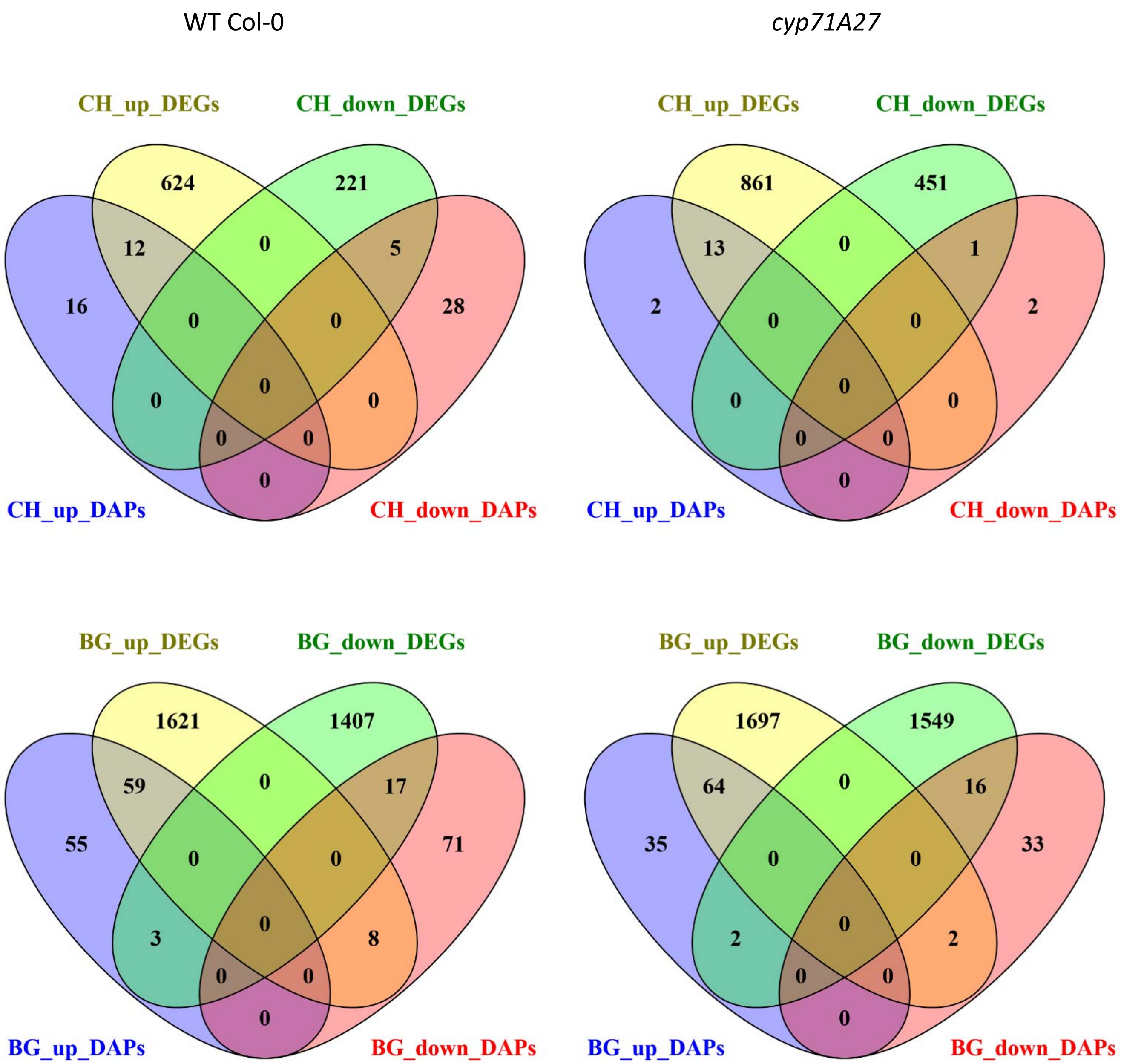


**Supplemental Figure S10.** Venn diagrams of DEGs and DAPs after treatment of WT Col-0 (left panels) and *cyp71A27* (right panels) with CH267 (CH) or *B. glumae* (BG).

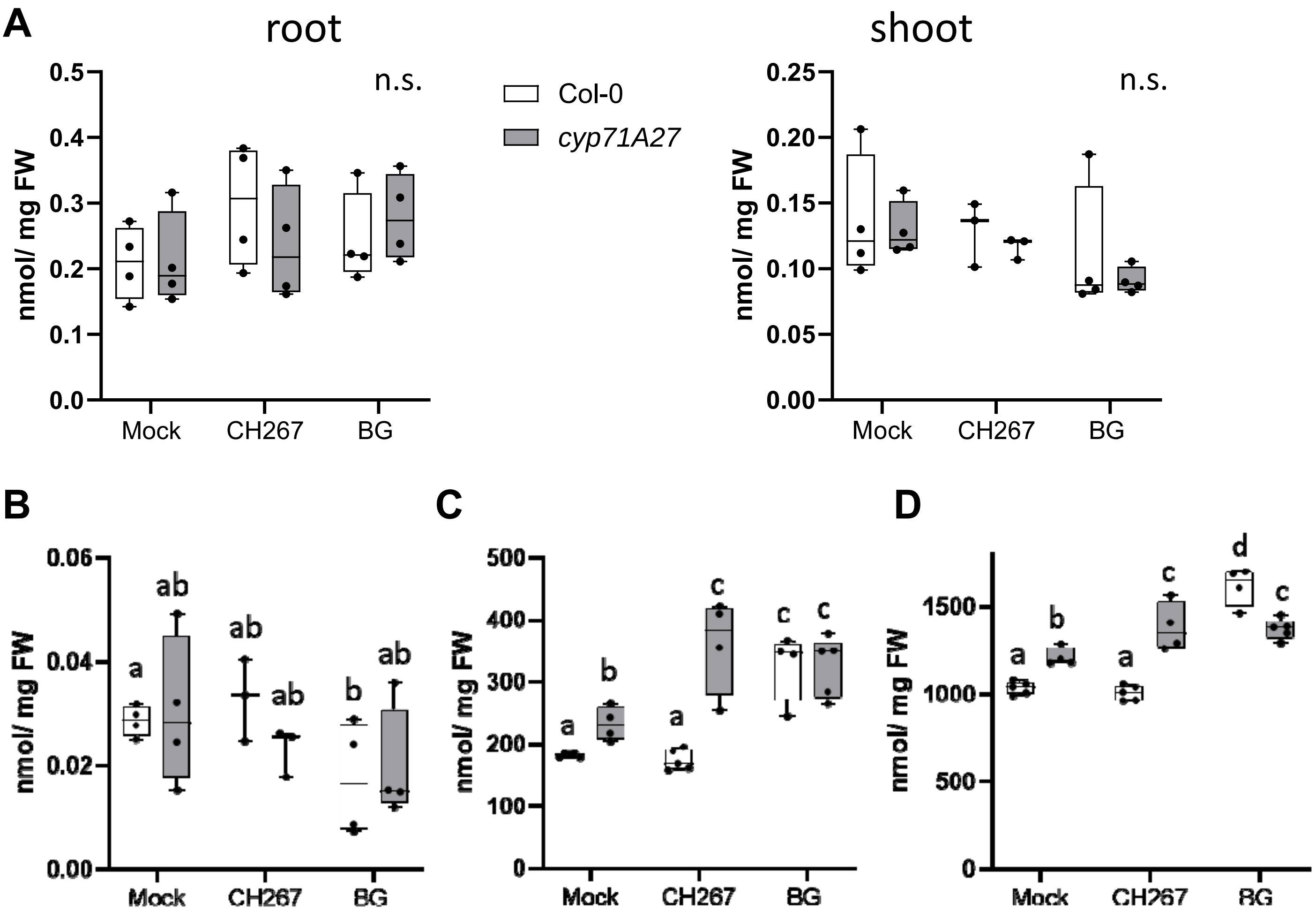


**Supplemental Figure S11.** Metabolite analysis of response of Arabidopsis to *Pseudomonas* sp. CH267 (CH) and *Burkholderia glumae* PG1 (BG). (**A**) aliphatic glucosinolates were determined in WT and *cyp71A27* roots and shoots treated with mock, CH, or BG by HPLC. (**B**) indolic glucosinolates (**C**) cysteine (**D**) glutathione in the shoots. Different letters mark values significantly different at p<0.05 (T-test). n.s. not significant.

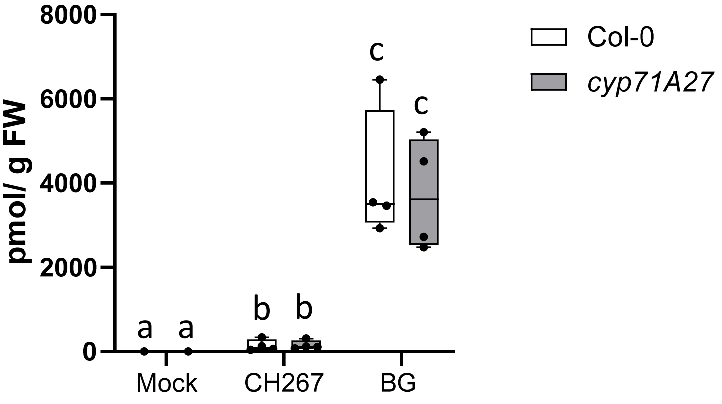


**Supplemental Figure S12.** Camalexin in exudates of Arabidopsis treated with *Pseudomonas* sp. CH267 and *Burkholderia glumae* PG1 (BG). Camalexin was determined in exudates of WT and *cyp71A27* treated with mock, CH267, or BG by HPLC. Different letters mark values significantly different at $p<0.05$ (T-test). n.s. not significant.

**A**

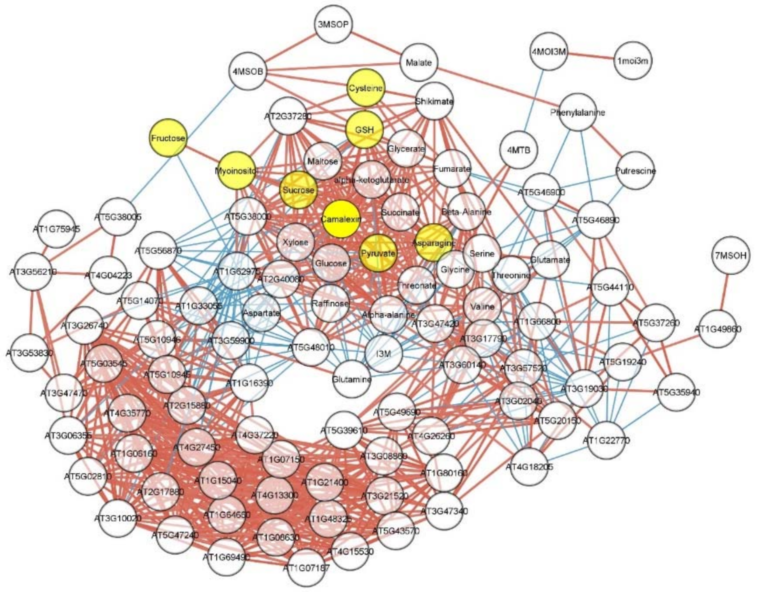

**B**

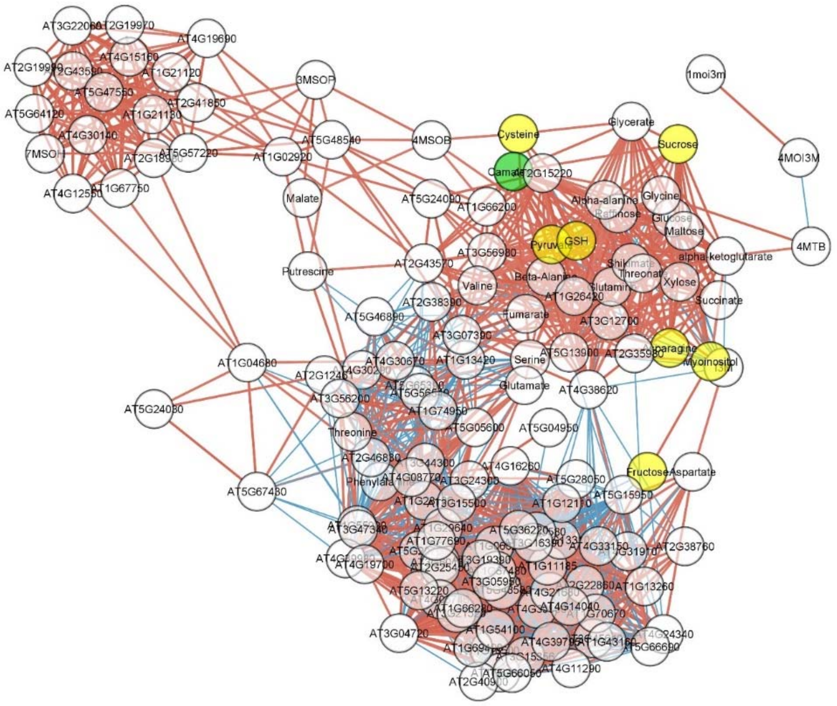

**C**

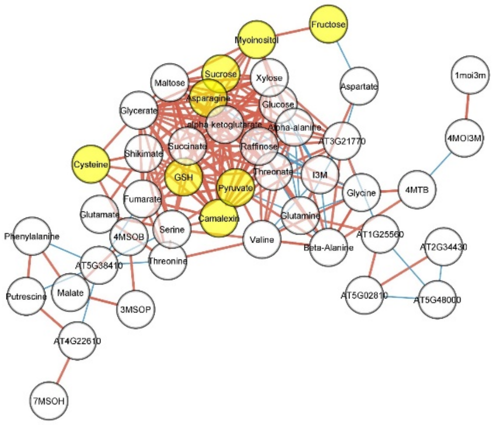


**Supplemental Figure S13.** Network analysis of DEGs and metabolites. The networks were created by pairwise correlations between DEGs and metabolite levels corresponding to (**A**) *cyp71A27* main effect (**B**) interaction cyp27 and CH267, and (**C**) interaction cyp27 and *B. glumae* The networks were visualized in Cytoscape; brown lines represent positive correlation and blue lines denote negative correlation. Yellow circles mark metabolites for construction of subnetworks in Figure 6.
